# H3.1K27me1 maintains transcriptional silencing and genome stability by preventing GCN5-mediated histone acetylation

**DOI:** 10.1101/2020.07.17.209098

**Authors:** Jie Dong, Chantal LeBlanc, Axel Poulet, Benoit Mermaz, Gonzalo Villarino, Kimberly M. Webb, Valentin Joly, Josefina Mendez, Philipp Voigt, Yannick Jacob

## Abstract

In plants, genome stability is maintained during DNA replication by the H3.1K27 methyltransferases ATXR5 and ATXR6, which catalyze the deposition of K27me1 on replication-dependent H3.1 variants. Loss of H3.1K27me1 in *atxr5 atxr6* double mutants leads to heterochromatin defects, including transcriptional de-repression and genomic instability, but the molecular mechanisms involved remain largely unknown. In this study, we identified the conserved histone acetyltransferase GCN5 as a mediator of transcriptional de-repression and genomic instability in the absence of H3.1K27me1. GCN5 is part of a SAGA-like complex in plants that requires ADA2b and CHR6 to mediate the heterochromatic defects of *atxr5 atxr6* mutants. Our results show that Arabidopsis GCN5 acetylates multiple lysine residues on H3.1 variants *in vitro,* but that H3.1K27 and H3.1K36 play key roles in inducing genomic instability in the absence of H3.1K27me1. Overall, this work reveals a key molecular role for H3.1K27me1 in maintaining genome stability by restricting histone acetylation in plants.

## Introduction

Genome and epigenome instability have been implicated in many human diseases, including cancer and neurodegenerative disorders. In proliferating cells, key mechanisms are required to properly copy DNA and different epigenetic states of the genome in the context of ongoing transcription and DNA repair. Chromatin replication is therefore a complex molecular operation that can lead to genomic rearrangements and other types of deleterious mutations in the absence of mechanisms preserving genome stability (1, 2).

Epigenetic information plays multiple regulatory roles during S phase of the cell cycle that are required to maintain genome stability in eukaryotes. In plants, one of the most well-studied genome maintenance pathways involves the histone post-translational modification (PTM) H3K27me1. The loss of H3K27me1 results in the loss of transcriptional silencing at heterochromatic loci and defects in the structural organization of heterochromatin (3, 4). In addition, decreased levels of H3K27me1 induce genome instability characterized by the presence of an excess of repetitive DNA (e.g. transposons) mainly in pericentromeric heterochromatin (hereafter referred to as heterochromatin amplification) (5). H3K27me1 is catalyzed by the plant-specific histone methyltransferases ATXR5 and ATXR6 (ATXR5/6), which are recruited to replication forks during DNA replication (3, 6, 7). Biochemical and structural studies have revealed that the SET domains of ATXR5/6 can methylate replication-dependent H3.1 variants, but not replication-independent H3.3 variants (8). These observations indicate that ATXR5/6 maintain H3K27me1 by methylating newly synthesized H3.1 variants (H3.1K27me1) during DNA replication, which protects against transcriptional de-repression and heterochromatin amplification. The precise molecular mechanism responsible for heterochromatin amplification in the absence of H3.1K27me1 remains unknown. However, a previous study suggested that transcriptional de-repression in the heterochromatin of *atxr5/6* double mutant plants is the cause of the genomic instability phenotype, potentially by inducing collisions between the transcription machinery and replication forks, and/or through R-loop formation (9). Based on this model, it is predicted that ATXR5/6-catalyzed H3.1K27me1 plays a key role in blocking transcriptional activity in the heterochromatin of plants.

Many PTMs on histones function as recruitment signals for chromatin “reader” proteins, which promote specific cellular activities like transcription at genomic regions enriched in these histone PTMs (10). Multiple studies have shown that methylation at H3K27 regulates transcriptional activity through various mechanisms, which are related to the specific methylation level (i.e., me1, me2 or me3) at K27. For example, H3K27me3 is involved in the recruitment of the repressive PRC1 complex in animals (11), and this role is conserved in plants (12). In contrast to H3K27me3, H3K27me1 and H3K27me2 are not as well characterized in animals, but they have specific effects on the regulation of transcriptional activity that do not appear to involve recruitment of chromatin readers. In mouse embryonic stem cells (ESCs), H3K27me2 is present on the majority of total histone H3 in chromatin and safeguards against unintended transcription by blocking CBP/p300-mediated H3K27 acetylation (H3K27ac) at non-cell-type-specific enhancers (13). In contrast, H3K27me1 is present at less than 5% of total H3s in ESCs, and is associated with transcriptionally active genes and contributes to their expression (13). However, the mechanism by which H3K27me1 performs this function remains unknown. Predicting the role of ATXR5/6-catalyzed H3K27me1 in plants based on comparative analysis with H3K27me1/me2 in animals is challenging, as it shares the same methylation level of transcriptionally-permissive H3K27me1, but its function in heterochromatin silencing in plants suggests properties related to H3K27me2. An additional similarity between plant H3K27me1 and animal H3K27me2 is that these histone PTMs are widely distributed and very abundant in their respective genomes. In Arabidopsis, H3K27me1 was estimated to be present on more than 50% of total H3 in inflorescence tissues (14), and it is enriched in transcriptionally silent regions of the genome (5). These observations suggest that H3.1K27me1 in plants may serve to block H3.1K27ac, providing a potential molecular mechanism for the role of ATXR5/6 in preventing transcriptional derepression and genomic instability in plants.

In this work, we identify the conserved histone acetyltransferase GCN5 as a mediator of transcriptional de-repression and heterochromatin amplification in the absence of H3.1K27me1 in plants. GCN5 cooperates with the transcriptional co-activator ADA2b and the chromatin remodeler CHR6 to induce these heterochromatic phenotypes. Our results also show that H3.1K36 plays a key role in inducing genome instability and transcriptional de-repression in the absence of H3.1K27me1, and that H3.1K27me1 interferes with GCN5-mediated acetylation at both H3.1K27 and H3.1K36. Overall, these results demonstrate the key role played by GCN5-mediated histone acetylation in contributing to the heterochromatin phenotypes observed in the absence of ATXR5 and ATXR6 in plants.

## Results

### Transcriptional de-repression and heterochromatin amplification in the absence of H3.1K27me1 are suppressed in *gcn5* mutants

One mechanism by which H3.1K27me1 might interfere with transcription in heterochromatin of plants is by preventing the deposition of H3.1K27ac, as methylation and acetylation at H3K27 have been shown to act antagonistically in other biological systems (15, 16). H3K27ac is catalyzed by multiple histone acetyltransferases in eukaryotes, including the widely conserved protein GCN5 (17–21). The Arabidopsis genome contains a single gene encoding a GCN5 ortholog (22). To assess if Arabidopsis GCN5 mediates the heterochromatin phenotypes associated with loss of H3.1K27me1, we created an *atxr5/6 gcn5* triple mutant by crossing a T-DNA insertion allele (SALK_030913) of *GCN5* into the hypomorphic *atxr5/6* mutant background (Supplemental Figure 1A) (3). Flow cytometry analyses showed strong suppression of heterochromatin amplification in the triple mutant as represented by the loss of the characteristic broad peaks corresponding to 8C and 16C endoreduplicated nuclei in *atxr5/6* mutants (Figure 1A). We also observed by microscopy that the heterochromatin decondensation phenotype of *atxr5/6* plants is suppressed in the *atxr5/6 gcn5* triple mutant (Figure 1B, Supplemental Figure 1B). A role for GCN5 in inducing genomic instability in *atxr5/6* was confirmed by using a different mutant allele of *gcn5* generated by temperature-optimized CRISPR/Cas9 (Supplemental Figure 1A, C and D) (23).

**Figure 1.**
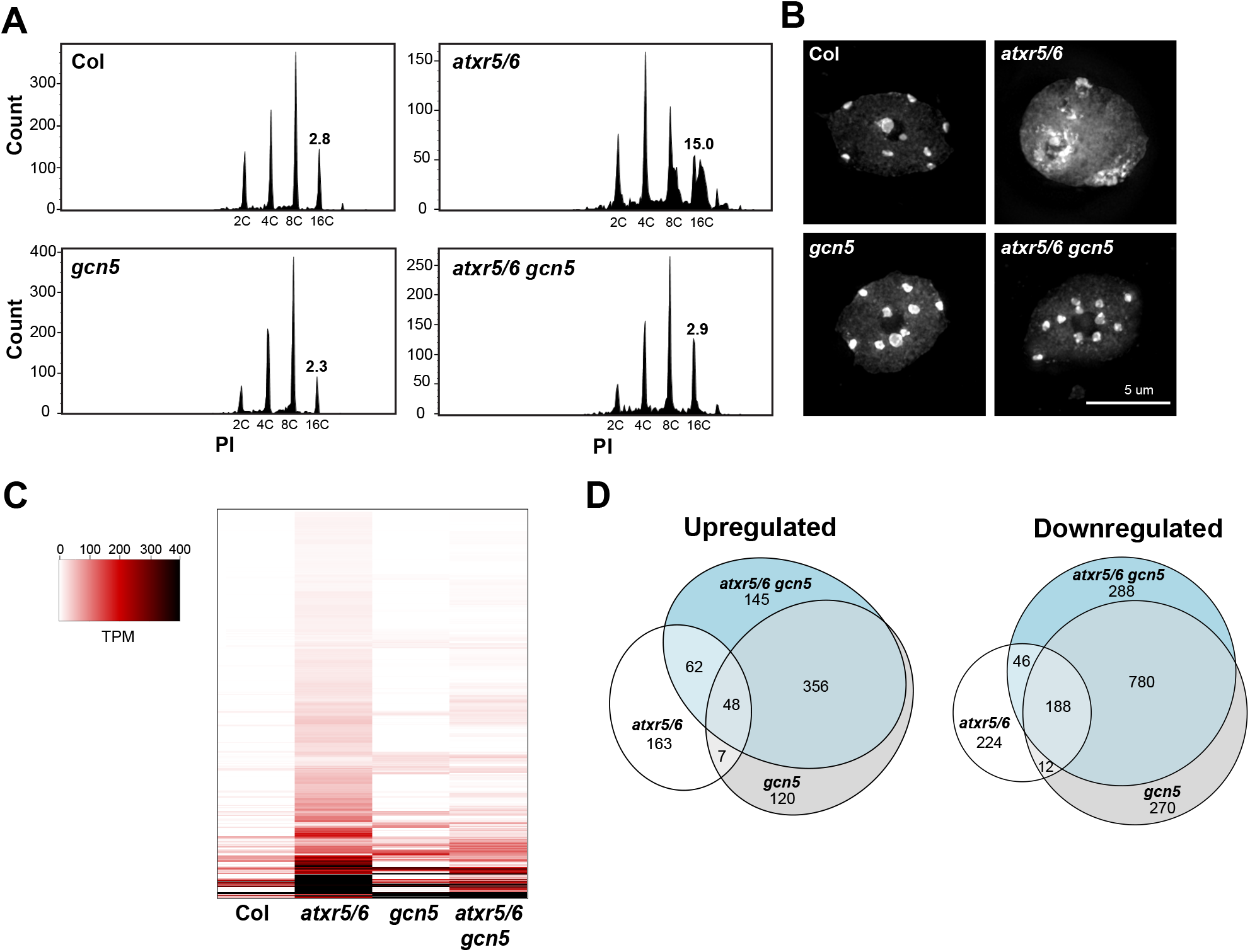
A mutation in *GCN5* suppresses transcriptional de-repression and heterochromatin amplification associated with H3.1K27me1 depletion. (A) Flow cytometry profiles of Col, *atxr5/6, gcn5* and *atxr5/6 gcn5* nuclei stained with propidium iodide (PI) with 2000 gated events. The numbers below the peaks indicate ploidy levels of the nuclei. The numbers above the 16C peaks indicate the robust coefficient of variation (CV). (B) Leaf interphase nuclei of Col, *atxr5/6, gcn5* and *atxr5/6gcn5* stained with DAPI. (C) Heat map showing the relative expression levels of *atxr5/6*-induced TEs (Supplemental Table 1) as measured by TPM (transcripts per million) in Col, *atxr5/6, gcn5* and *atxr5/6 gcn5*. (D) Euler diagrams showing the upregulated and downregulated genes (2-fold change) in *atxr5/6, gcn5* and *atxr5/6 gcn5* in comparison to Col plants *(Padj* < 0.05).

To measure the impact of GCN5 on transcriptional de-repression in *atxr5/6* mutants, we performed RNA-seq analyses and observed widespread suppression of transposable element (TE) reactivation in the *atxr5/6 gcn5* triple mutants compared to *atxr5/6*, although some TEs remained de-repressed compared to Col (Figure 1C and Supplemental Table 1). Although *GCN5* has a genome-wide impact on transcription as shown by the 1781 misregulated genes in *gcn5* single mutants (Figure 1D, Supplemental Table 2), none of the known transcriptional suppressors of *atxr5/6* mutants *(SERRATE [SE], AtTHP1, AtSAC3B, AtSTUbL2, AtMBD9* and *DDM1*) are downregulated in *gcn5* mutants or *atxr5/6 gcn5* triple mutants (Supplemental Figure 1E) (9, 24), indicating that suppression of the heterochromatin phenotypes in *atxr5/6 gcn5* is not the result of decreased expression levels of these genes.

### GCN5 functions with ADA2b and CHR6 to disrupt heterochromatin in the absence of H3.1K27me1

GCN5 is a member of the multi-subunit SAGA complex which acts as a transcriptional coactivator in yeast and animals in part by modifying chromatin (25). Key components of this complex are the proteins GCN5, ADA2, ADA3 and SGF29, which form the histone acetylation module of SAGA (Figure 2A). The Arabidopsis genome contains single genes encoding GCN5 and ADA3 and two genes each encoding ADA2 *(ADA2a* and *ADA2b)* and SGF29 *(SGF29a* and *SGF29b)* (26). *gcn5* and *ada2b* single mutants show pleiotropic phenotypes, which are also shared by the *atxr5/6 gcn5* and *atxr5/6 ada2b* mutants, respectively (Supplemental Figure 2A) (27). To test if ADA2b is also required for inducing the heterochromatin phenotypes of *atxr5/6* mutants, we created an *atxr5/6 ada2b* triple mutant and observed, similarly to *atxr5/6 gcn5* mutants, that genomic instability is suppressed in that background (Figure 2B). This result is supported by altered expression of *BRCA1,* which functions in eukaryotes as a DNA-damage response gene involved in maintaining genome stability (28, 29). As previously reported, *BRCA1* levels are upregulated in *atxr5/6* (4), and our results show that both *ADA2b* and *GCN5* are required for this induction (Figure 2C and Supplemental Figure 2B). Similarly to *gcn5,* an *ada2b* mutation in *atxr5/6* suppresses transcriptional de-repression of the heterochromatic *TSI* DNA repeat (Figure 2D and Supplemental Figure 2C).

**Figure 2.**
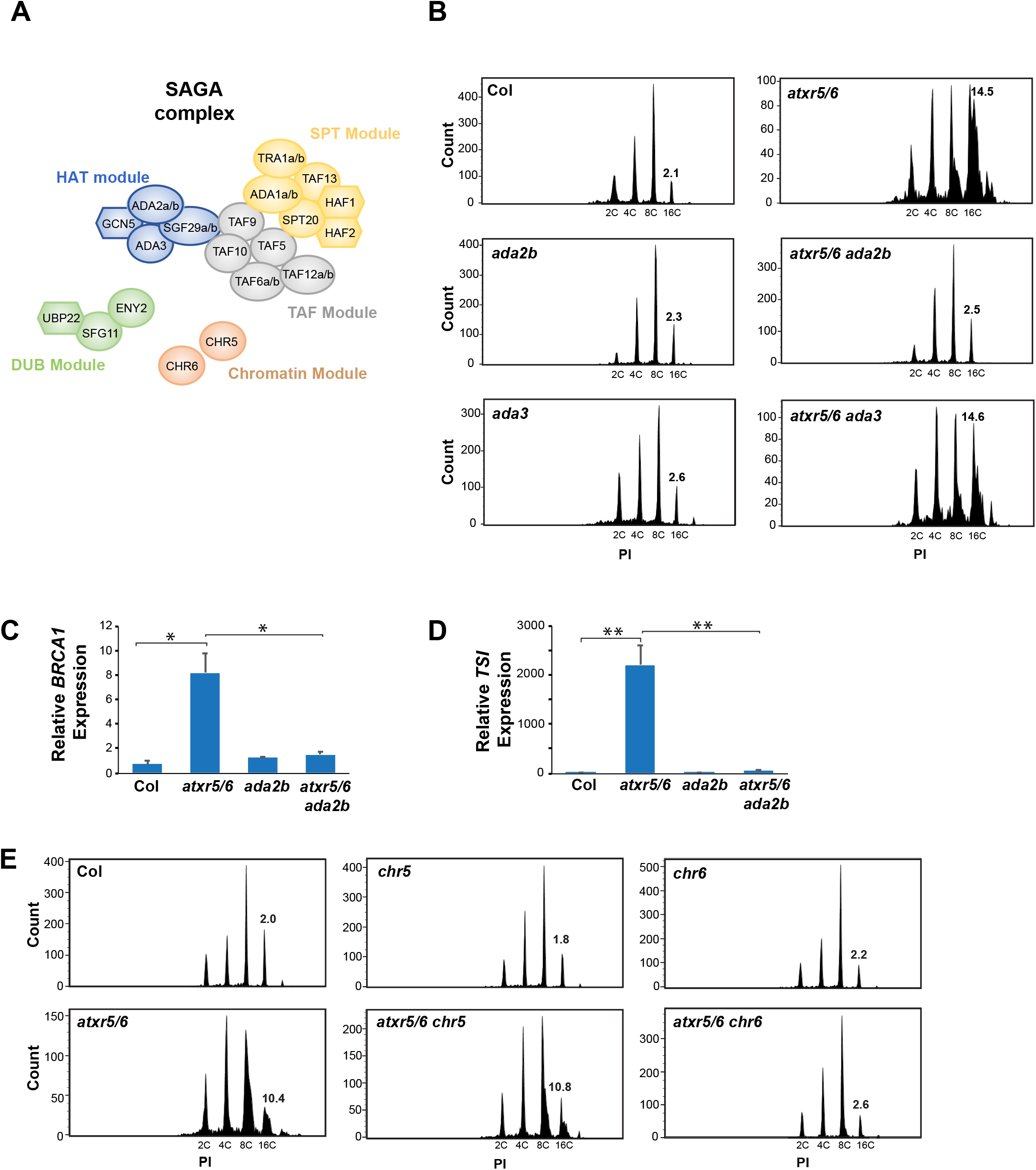
GCN5 requires ADA2b to induce heterochromatic defects in *atxr5/6* mutants. (A) Proposed subunits of the Arabidopsis SAGA complex. HAT: histone acetylation module; DUB: deubiquitination module; SPT: recruiting module; TAF; coactivator architecture module. (B) Flow cytometry profiles of Col, *atxr5/6, ada2b, atxr5/6 ada2b, ada3,* and *atxr5/6 ada3.* (C and D) RT-qPCR analyses of *BRCA1* (C) and the repetitive element *TSI* (D) in Col, *atxr5/6, ada2b* and *atxr5/6 ada2b.* Data represent the mean of three biological replicates and error bars indicate the standard error of the mean (SEM). Unpaired t-test: * *p* < 0.01, ** *p* < 0.001. (E) Flow cytometry profiles of Col, *atxr5/6, chr5, atxr5/6 chr5, chr6,* and *atxr5/6 chr6.*

Next, we generated an *atxr5/6 ada3* triple mutant, but unlike *atxr5/6 ada2b,* it did not suppress the genome instability phenotype associated with the *atxr5/6* double mutant (Figure 2B). The reported ADA3 protein in Arabidopsis displays low similarity to the ADA3 orthologs from yeast and human (26.3% and 16.3%, respectively, compared to >35% similarity for GCN5 and ADA2b (30)), and might therefore have diverged and not be required for GCN5 and ADA2b to acetylate histones in plants. To further investigate whether other modules of SAGA mediate the heterochromatin phenotypes associated with the loss of H3.1K27me1, we created triple mutant combinations between *atxr5/6* and mutated alleles of genes proposed to encode subunits of the deubiquitination module of SAGA in plants (Figure 2A). Our results show that the Arabidopsis orthologs of ENY2, UBP22 and SGF11 are not required for inducing heterochromatin amplification, indicating that the heterochromatin phenotypes of *atxr5/6* are independent of the deubiquitination function of SAGA (Supplemental Figure 2D). Finally, we created triple mutant combinations between *atxr5/6* and *chr5* or *chr6.* CHR5 and CHR6 are both chromatin remodeling enzymes that have been proposed to be present in the SAGA complex in plants (Figure 2A). CHR5 is the most closely related plant protein to CHD1-type chromatin remodelers that are part of the SAGA complex in yeast and mammals (26, 30), while CHR6 (also known as CHD3/PICKLE) has been shown to co-purify with SAGA subunits from Arabidopsis tissue (31). Our results show that heterochromatin amplification is suppressed in the *atxr5/6 chr6* triple mutant, but not *atxr5/6 chr5* (Figure 2E). Overall, these results indicate essential roles for ADA2b and CHR6 in mediating the heterochromatic phenotypes observed in the absence of H3.1K27me1.

### GCN5-mediated H3.1K27ac induces the heterochromatin defects associated with loss of H3.1K27me1

The GCN5 orthologs in yeast and mammals have been shown to acetylate multiple lysine residues of histone H3 (i.e., K9, K14, K18, K23, K27 and K36) *in vitro,* however, substrate specificity in the context of different histone H3 variants has never been tested for any GCN5 ortholog (18, 20). In addition, while the Arabidopsis GCN5 has been shown to acetylate H3K9 and H3K14 on H3 peptides *in vitro* (32), acetylation at H3K27 by the Arabidopsis GCN5 ortholog has not been tested.

To investigate the substrate specificity of GCN5, we performed *in vitro* histone lysine acetyltransferase (HAT) assays using recombinant nucleosomes containing either plant histone H3.1 or H3.3 variants. We recombinantly expressed and purified an Arabidopsis protein complex composed of GCN5 and ADA2b (Supplemental Figure 3). Our results show that GCN5 has HAT activity at K9, K14, K18, K23, K27 and K36 of histone H3 (Figure 3A). In contrast to ATXR5/6, the enzymatic activity of GCN5 at H3K27 is not regulated by H3 variants, as H3.1 and H3.3 nucleosomes show equivalent acetylation levels in our HAT assays (Figure 3A). As controls for these results, we used H3.1K27ac and H3.3K27ac peptides to validate that the H3K27ac antibody used did not show preference for H3.1 or H3.3 (Figure 3B), and we validated the specificity of this antibody by using H3K27M nucleosomes (Figure 3C). Similarly to H3K27, we did not observe any major difference in histone acetyltransferase activity between H3.1 and H3.3 nucleosomes at the other lysine substrates of Arabidopsis GCN5 (Figure 3A). We also confirmed that H3.1K27me1 prevents acetylation by GCN5 at K27 by using recombinant nucleosomes mono-methylated at K27 (Figure 3D). To assess if H3.1K27ac mediates the heterochromatin phenotypes present in *atxr5/6* mutants *in vivo,* we introduced into wild-type plants an H3.1K27Q transgene. Replacement of lysine (K) with glutamine (Q) in histones has been used for *in vivo* chromatin studies to partially mimic the acetylated state of histone lysine residues (33–35). Our analyses of first-generation transformed (T1) plants show that expression of H3.1K27Q in wild-type plants is sufficient to induce transcriptional de-repression of the heterochromatic *TSI* repeat (Figure 3E) and activation of the genome instability marker *BRCA1* (Figure 3F), which are both specifically upregulated in *atxr5/6* mutants. Overall, these results support a specific role for GCN5-mediated H3.1K27ac in inducing the heterochromatic phenotypes associated with loss of H3.1K27me1 in *atxr5/6* mutants.

**Figure 3.**
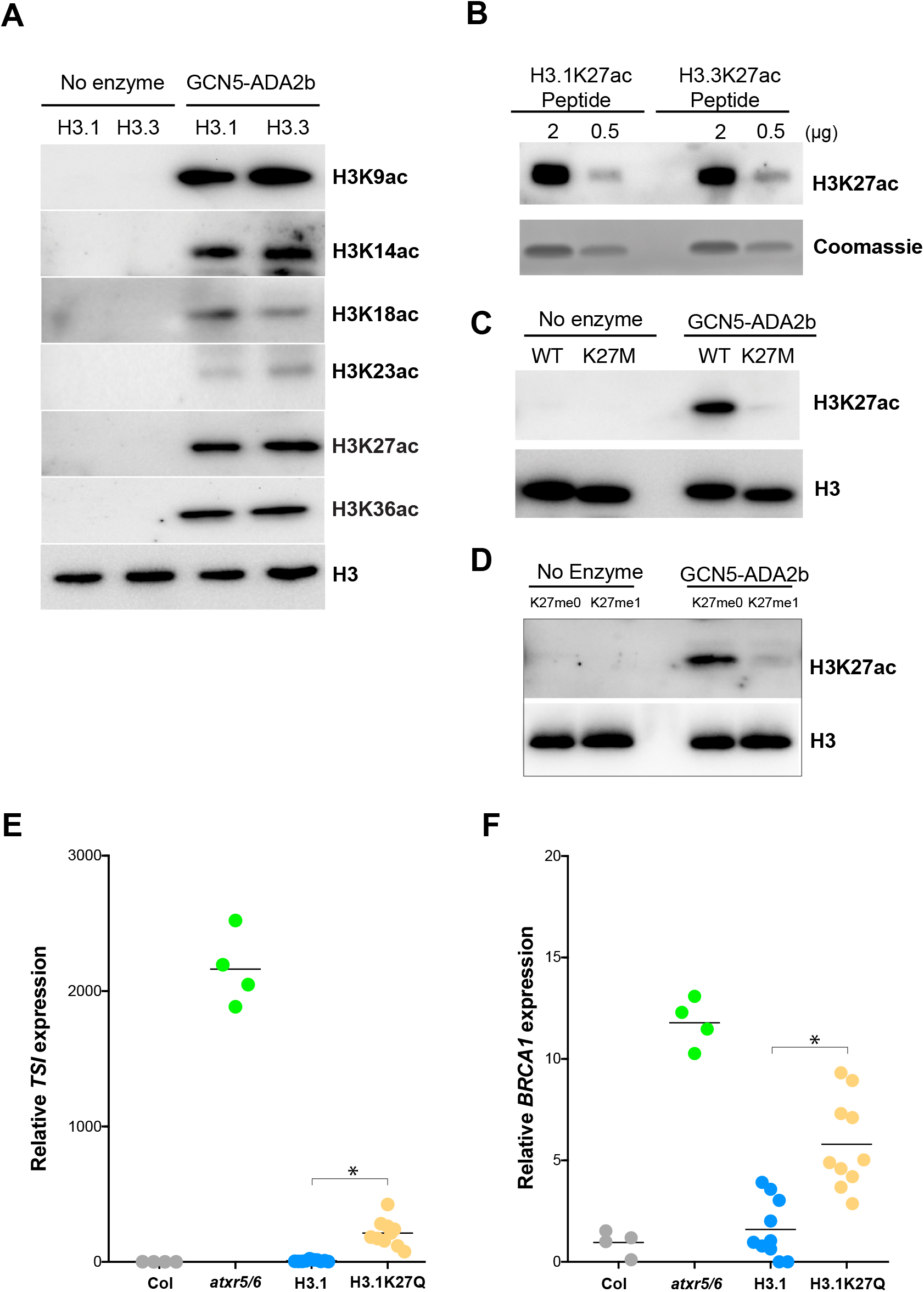
Arabidopsis GCN5 acetylates H3.1K27 and induces the heterochromatic defects associated with *atxr5/6.* (A) *In vitro* HAT assays with the GCN5-ADA2b complex and H3.1 and H3.3 nucleosomes using anti-H3K9ac, anti-H3K14ac, anti-H3K18ac, anti-H3K23ac, anti-H3K27ac, anti-H3K36ac and anti-H3 antibodies. (B) Western blot of H3.1K27ac and H3.3K27ac peptides with H3K27ac antibody. (C) *In vitro* HAT assay with the GCN5-ADA2b complex and H3K27M nucleosomes. (D) *In vitro* HAT assays with the GCN5-ADA2b complex and H3K27me0 and H3K27me1 nucleosomes using anti-H3K27ac, anti-H3 antibodies (E and F) RT-qPCR for the heterochromatic transcriptional reactivation marker *TSI* (E) and the genome stability marker *BRCA1* (F) in Col, *atxr5/6* and first-generation transformed (T1) plants expressing WT H3.1 or H3.1K27Q. Ten independent T1 plants were used in the experiments. Unpaired t-test: * *p* < 0.001.

### H3.1K36 is required to induce genome instability in the absence of H3.1K27me1

Our *in vitro* results suggest that, in addition to K27, other lysine residues on H3.1 could contribute to GCN5-mediated genomic instability in the absence of H3.1K27me1. To assess this, we set up a suppressor screen based on *in vivo* replacement of histone H3.1 with the point mutant H3.1S28A. Replacement of serine with alanine on H3.1 variants at position 28 (H3.1S28A) generates H3.1 substrates that cannot be methylated by ATXR5/6 (Figure 4A) (36). In contrast, H3.1S28A can still be methylated at K27 by plant PRC2-type complexes and acetylated by the GCN5-ADA2b complex, albeit at lower efficiencies (Supplemental Figure 4A-B). We transformed the H3.1S28A transgene into a mutant Arabidopsis background expressing a reduced amount of endogenous histone H3.1 (i.e., *h3.1* quadruple mutant (8)) and observed in T1 plants the phenotypes associated with loss of H3.1K27me1, including genomic instability as detected by flow cytometry and increased levels of the genome instability marker gene *BRCA1* (Figure 4B-C), and transcriptional de-repression of the heterochromatic *TSI* DNA repeat (Figure 4D). These results indicate that expression of H3.1S28A in plants generates phenotypes similar to *atxr5/6* mutants due to loss of H3.1K27me1. We then introduced a series of H3.1S28A expression constructs containing a second mutation (Lys to Arg replacement) at a residue known to be acetylated by GCN5 into the *h3.1* quadruple mutant, and T1 plants were assessed for the phenotypes associated with loss of H3.1K27me1. This targeted screen identified H3.1K36 as being essential for inducing genome instability, as flow cytometry analyses demonstrated that H3.1S28A K36R suppresses heterochromatin amplification, while the other targeted mutations do not (Figure 4B). The H3.1S28A K36R replacement line also rescued the increased expression of *BRCA1* (Figure 4C) and transcriptional de-repression of *TSI* (Figure 4D).

**Figure 4.**
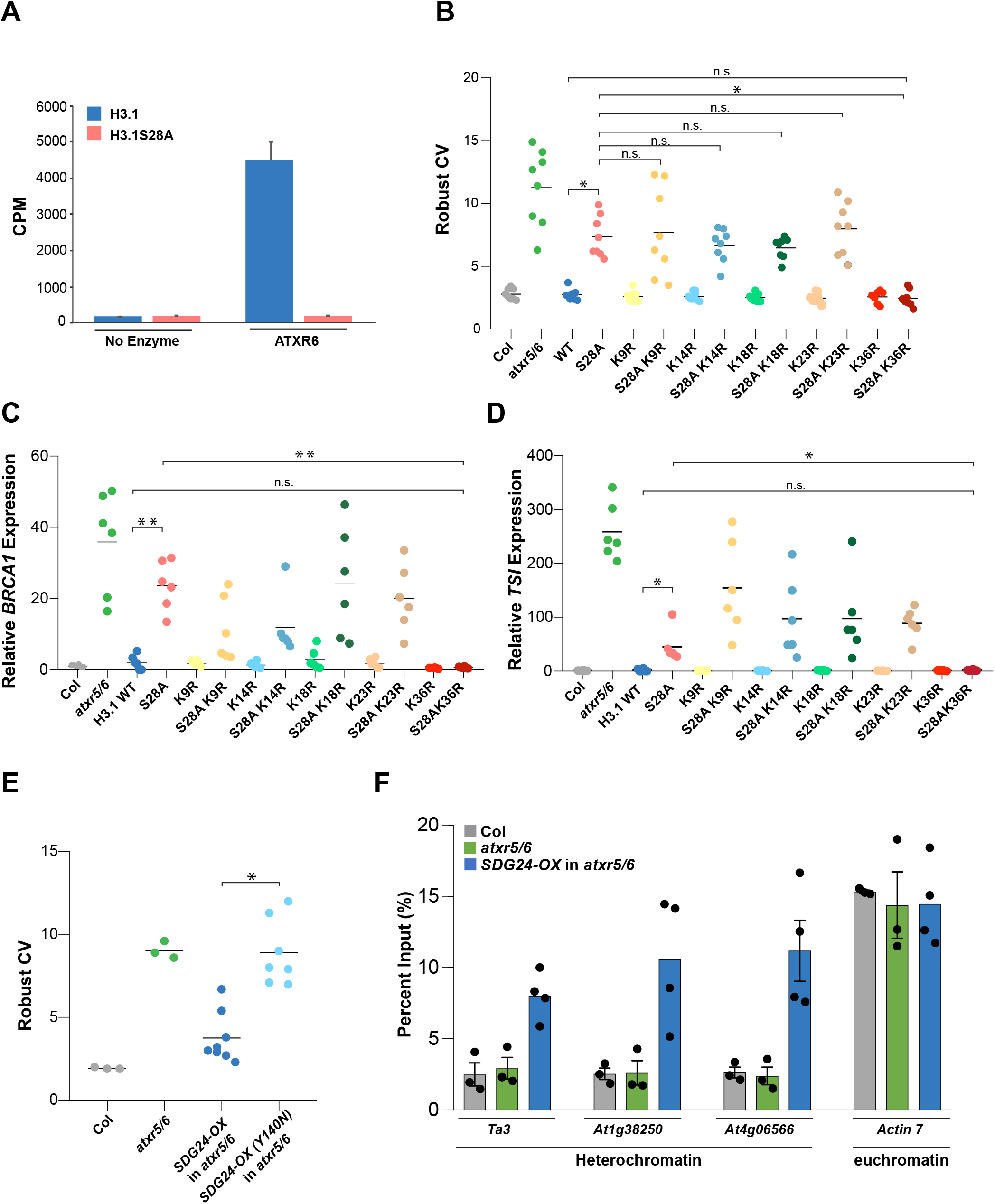
Heterochromatin amplification in the absence of H3.1K27me1 requires H3.1K36. (A) *In vitro* histone lysine methylation assays using H3.1 and H3.1S28A peptide substrates and ATXR6. The average of three experiments and SEM are shown. CPM; counts per minute (B) Robust CV values for 16N nuclei obtained by flow cytometry analysis. For Col and *atxr5/6*, each dot represents an independent biological replicate. For the H3.1 replacement lines, each dot represents one T1 plant. Horizontal bars indicate the mean. Unpaired t-test: * *p* < 0.0001 and n.s. = not significantly different. (C and D) RT-qPCR analyses of *BRCA1* (C) and the repetitive element *TSI* (D) in Col, *atxr5/6* and H3.1 replacement lines. For Col and *atxr5/6,* each dot represents an independent biological replicate. For the H3.1 lines, each dot represents one T1 plant. Horizontal bars indicate the mean. Unpaired t-test: \**p* < 0.01, \*\**p* < 0.0005 (E) Flow cytometry analyses showing robust CV values for 16N nuclei. For the *SDG24-OX* lines, each dot represents one T1 plant. Horizontal bars indicate the mean. Unpaired t-test: * *p* < 0.001. (F) H3K36me3 ChIP-qPCR at *Ta3, At1g38250, At4g06566* and *Actin7.* For Col and *atxr5/6*, each dot represents an independent biological replicate. For the *SDG24-OX* lines, each dot represents one T1 plant. Bars indicate the mean. Error bars indicate SEM.

Furthermore, expression of the H3.1S28A K36R mutant did not generate a serrated leaves phenotype as seen in all the other H3.1S28A lines (Supplemental Figure 5). As mutations at K9, K14, K18 and K23 on the H3.1 variant did not suppress the phenotypes associated with the H3.1S28A mutation, these results indicate a specific role for H3.1K36 in inducing genome instability in the absence of H3.1K27me1.

GCN5-mediated acetylation of H3.1K36 could be required to induce the heterochromatin defects of *atxr5/6* mutants. One prediction from this model is that increasing histone methylation at H3.1K36 (H3.1K36me) would result in the suppression of the *atxr5/6* mutant phenotypes, as H3.1K36me would antagonize H3.1K36 acetylation by GCN5. To test this, we constitutively expressed (using the 35S promoter) all five Arabidopsis H3K36 methyltransferases *(SDG4, SDG7, SDG8, SDG24,* and *SDG26)* in *atxr5/6* mutants (37, 38). We performed flow cytometry analyses on T1 plants and found that overexpression of *SDG24 (SDG24-OX)* strongly suppresses the heterochromatin amplification phenotype (Figure 4E). We did not observe a similar effect in T1 lines overexpressing *SDG4*, *SDG7*, *SDG8* or *SDG26* (Supplemental Figure 6). The ability of *SDG24-OX* to suppress heterochromatin amplification is dependent on SDG24 having a functional methyltransferase (SET) domain, as overexpression of an *SDG24* variant containing a point mutation (Y140N) in a conserved residue essential for SET domain activity does not suppress the phenotype (Figure 4E) (5, 39). We performed ChIP-qPCR experiments with *SDG24-OX* plants and detected an increase in H3K36me3 levels at heterochromatic regions *(Ta3, At1G38250, At4G06566)* known to be transcriptionally de-repressed in *atxr5/6* mutants (Figure 3F). Taken together, these results suggest that H3.1K36ac is required to induce transcriptional de-repression and heterochromatin amplification in the absence of H3.1K27me1.

### Loss of H3.1K27me1 in plants increases H3K27ac and H3K36ac deposition in heterochromatin

Our results support a model in which GCN5 acetylates both H3.1K27 and H3.1K36 in the absence of H3.1K27me1 to induce the heterochromatin phenotypes of *atxr5/6* mutants. To assess if H3.1K27me 1 depletion leads to an increase of H3K27ac and H3K36ac *in vivo,* we performed ChIP-seq with reference exogenous genome (ChIP-Rx) for H3K27ac and H3K36ac in Col (WT), *atxr5/6, gcn5* and *atxr5/6 gcn5* (40). We found that both histone marks are enriched at the 5’ end of protein-coding genes after the transcriptional start site (TSS) in Arabidopsis (Figure 5A), and this spatial distribution correlates with transcriptional activity, albeit not in a linear relationship (Supplemental Figure 7) (41, 42). Comparative analysis of H3K27ac and H3K36ac in Col and *gcn5* single mutants demonstrates that loss of GCN5 results in a decrease of H3K27ac and H3K36ac at euchromatic genes (Figure 5A).

**Figure 5.**
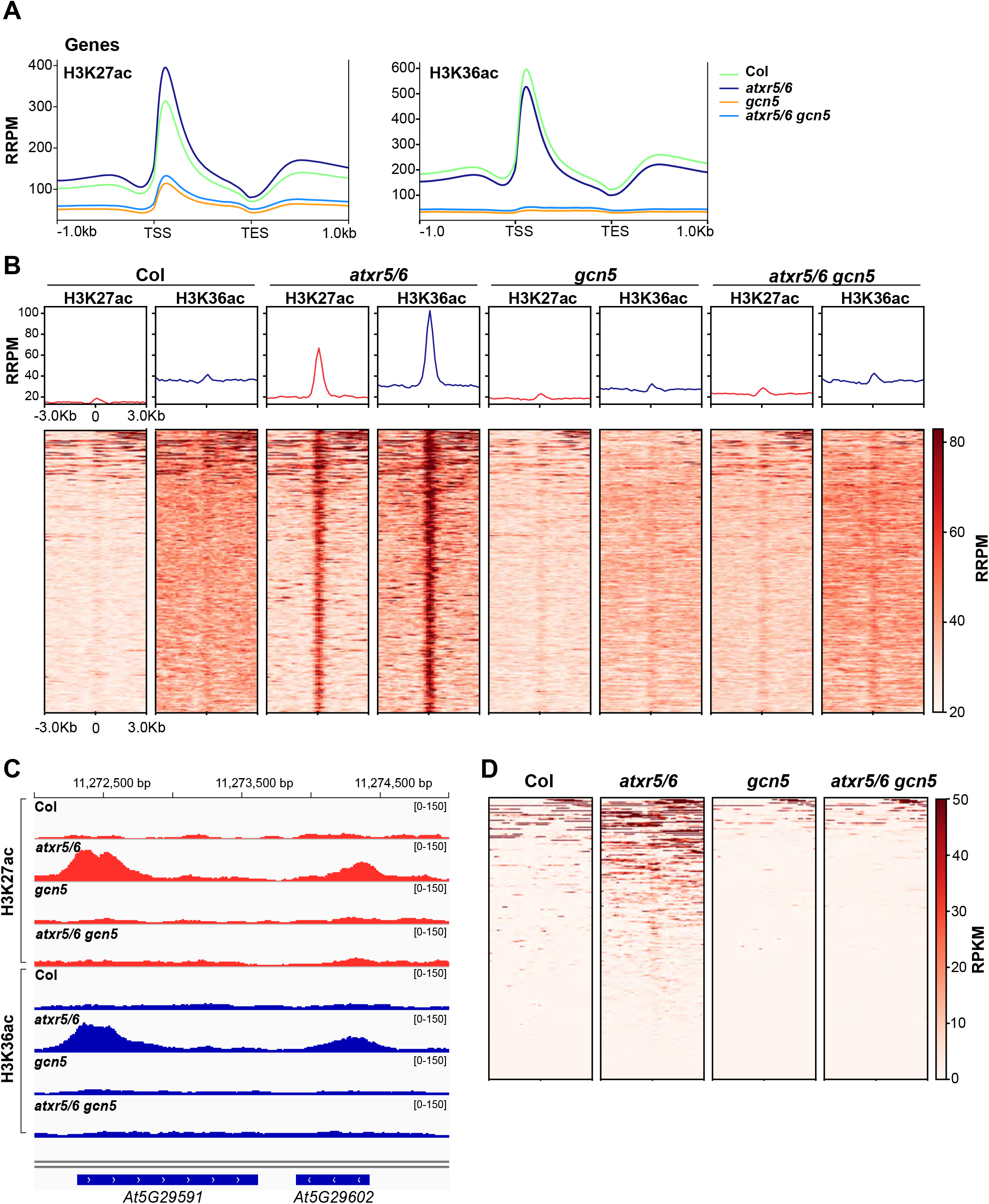
Mutations in *atxr5/6* lead to an increase in H3K27ac and H3K36ac in heterochromatin. (A) Normalized average distribution of H3K27ac and H3K36ac over protein coding genes for Col, *atxr5/6*, *gcn5* and *atxr5/6 gcn5* in reference-adjusted reads per million (RRPM). TSS, transcription start site; TES, transcription end site. (B) Normalized average distribution and heatmap of H3K27ac and H3K36ac normalized reads surrounding the H3K27ac/H3K36ac enriched heterochromatic regions in *atxr5/6* compared to Col. (C) Genome browser snapshot showing normalized H3K27ac and H3K36ac ChIP-seq data over a region of chromosome 5 that includes TE genes *At5g29591* and *At5g29602.* The y-axis unit is RRPM. (D) Heatmap showing the RNA-seq reads mapping to the region 3 kb around the center of H3K27ac/H3K36ac peaks as measured by RPKM (reads per kilobase million) in Col, *atxr5/6*, *gcn5* and *atxr5/6 gcn5*.

Focusing on heterochromatin, which we defined based on previously identified chromatin states in Arabidopsis (Supplemental Table 3) (43), we identified multiple regions that were enriched in both H3K27ac and H3K36ac in *atxr5/6* but not in Col plants (Figure 5B-C and Supplemental Table 4). H3K27ac and H3K36ac enrichment in heterochromatin was greatly decreased in *atxr5/6gcn5* triple mutants (Figure 5B-C), suggesting that higher levels of H3K27ac and H3K36ac in heterochromatic regions of *atxr5/6* are almost completely dependent on GCN5. We next tested if the de-repressed TEs in *atxr5/6* identified in by RNA-seq overlap or are in close proximity (+/-3kb) to the genomic regions showing increased levels of H3K27ac and H3K36ac in *atxr5/6.* We observed a large overlap between transcriptionally de-repressed genomic regions and regions enriched in H3K27ac and H3K36ac in *atxr5/6* mutants (Figure 5D). The regions shown in Figure 5D likely represent a low estimate of the total overlap between H3K27ac/H3K36ac regions and transposon reactivation due to the inherent lack of sensitivity of ChIP-seq and RNA-seq experiments in backgrounds showing low-level TE de-repression, like *atxr5/6* mutants. For example, we found that a 5-fold increase in sequencing depth (75 versus 15 million reads) in our RNA-seq experiments resulted in a 43% increase in the number of de-repressed TEs identified in *atxr5/6* (446 TEs versus 312 TEs) (Supplemental Table 1). To further demonstrate the sensitivity issue associated with low-level de-repression in *atxr5/6,* we performed RT-qPCR on multiple TEs that showed an increase in H3K27ac in *atxr5/6,* but were not identified as differently expressed by RNA-seq. For many of these TEs, including *At1g36040* and *At5g29602* (Supplemental Figure 8), we observed higher expression levels in *atxr5/6* compared to wild-type plants, thus confirming the limitations of genome-wide sequencing for detecting low-level TE de-repression in *atxr5/6* mutants. Taken together, these results demonstrate that the loss of H3.1K27me1 in *atxr5/6* mutants leads to GCN5-dependent increase of H3K27ac and H3K36ac in heterochromatin.

### H3.1K27me1 regulates the deposition of H3.1K36ac by GCN5

Methylation and acetylation at H3K27 have an antagonistic relationship in the genome of animals, which is mediated by the interplay between the H3K27 methyltransferase complex PRC2 (H3K27me) and the histone acetyltransferases p300 and CBP responsible for H3K27ac (15, 16). Our work supports a similar relationship in plants at K27 on H3.1 variants that is mediated by different enzymes, with ATXR5/6-catalyzed H3.1K27me1 preventing the acetylation of H3.1K27 by GCN5. Interactions between post-translational modifications on different histone residues also contribute to chromatin regulation in eukaryotes. One example of this is the inhibition of PRC2 activity towards H3K27 when H3K36 is di- or trimethylated on the same histone (44–46). This suggests that the activity of other chromatin-modifying enzymes may be affected by crosstalk between modified forms of H3K27 and H3K36. To assess if acetylation of H3.1K36 by GCN5 is regulated by H3.1K27me1, we performed *in vitro* HAT assays using recombinant plant nucleosomes containing either unmodified H3.1 or H3.1K27me1. In these assays, we consistently observed a 40% decrease in the levels of acetylation at H3.1K36 on nucleosomes mono-methylated at H3.1K27 compared to unmodified H3.1 (Figure 6A-B). This effect of H3.1K27me1 on Arabidopsis GCN5 activity appears to be specific to H3.1K36, as GCN5-mediated acetylation of H3.1K9 was not affected by mono-methylation at K27. We also tested if reciprocally, methylation at H3.1K36 would affect acetylation at K27 by GCN5. We did not observe any difference in acetylation levels at K27 using K36me0 and K36me3 nucleosomes (Figure 6C). Overall, these results suggest that ATXR5/6-catalyzed H3.1K27me1 in plants interferes with GCN5-mediated acetylation at both H3.1K27 and H3.1K36.

**Figure 6.**
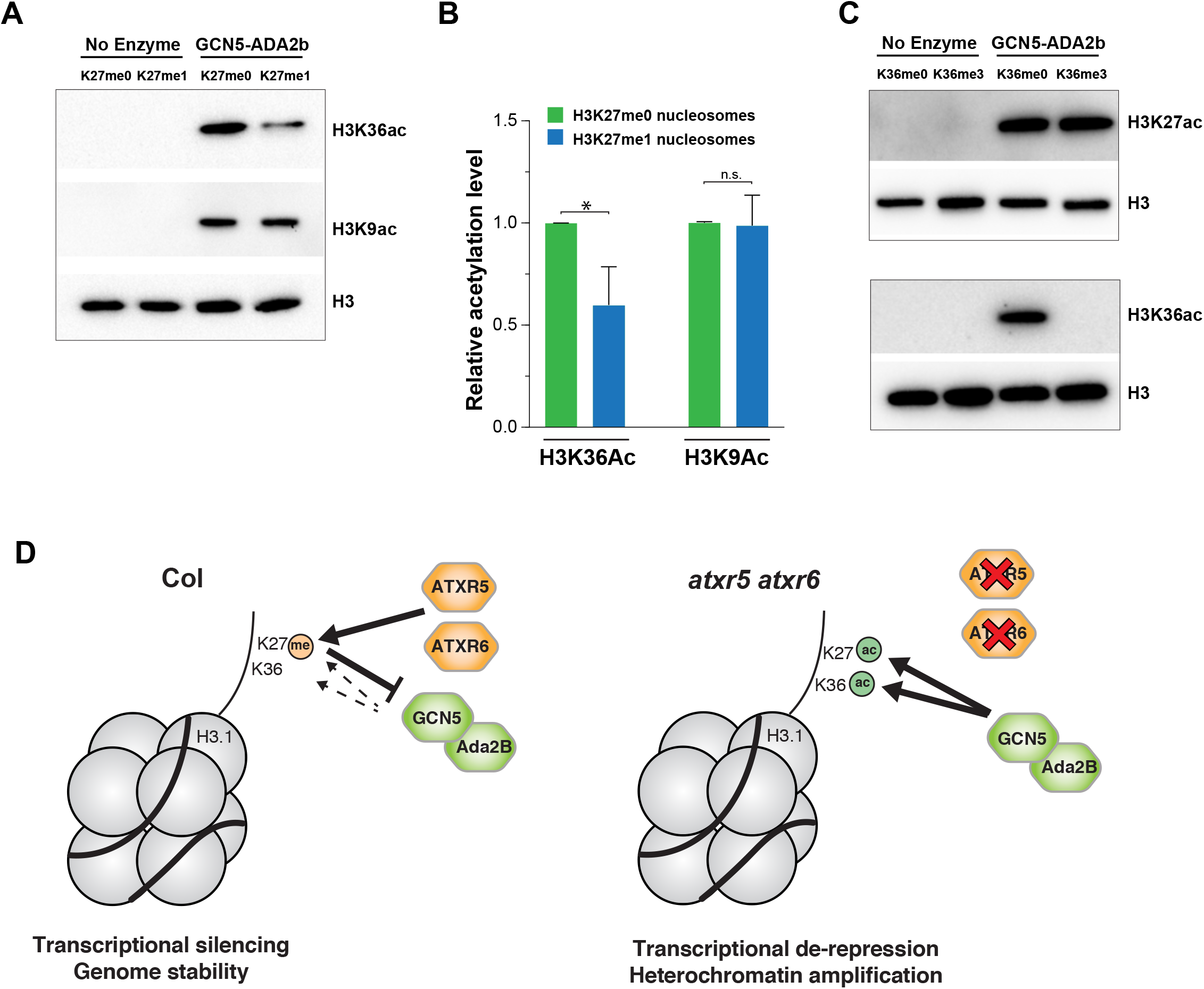
H3K36 acetylation by the GCN5-ADA2b complex is regulated by H3K27me1. (A) *In vitro* HAT assays with the GCN5-ADA2b complex and H3K27me0 and H3K27me1 nucleosomes. (B) Data from three technical replicates of HAT assays with the GCN5-ADA2b complex and H3K27me0 and H3K27me1 nucleosomes. Unpaired t-test: * *p* < 0.05, and n.s. = not significantly different. (C) *In vitro* HAT assays with the GCN5-ADA2b complex and H3K36me0 and H3K36me3 nucleosomes. D) Model depicting the role of H3.1K27me1 in blocking GCN5-mediated acetylation of H3.1K27ac and H3.1K36ac.

## Discussion

Previous work had suggested that transcriptional reactivation of heterochromatic regions is responsible for inducing genomic instability in the absence of H3.1K27me1 in plants (9). However, the mechanism by which H3.1K27me1 prevents transcriptional de-repression in heterochromatin had remained unknown. Our study supports a model where ATXR5/6-mediated H3.1K27me1 serves to prevent a SAGA-like complex that includes GCN5, ADA2b and CHR6 from acetylating the H3.1 variant and initiating transcriptional de-repression (Figure 6D). K27me1 is the most abundant post-translational modification on H3.1K27 in plants (14), and our results suggest that it plays a role analogous to the one proposed for PRC2-catalyzed H3K27me2 in animals, which is present on 50-70% of total histone H3 in mouse embryonic stem cells, blocks H3K27ac, and prevents spurious transcription (13, 47, 48). In animals, p300 and CBP are the main histone acetyltransferases contributing to H3K27ac in the absence of PRC2-mediated H3K27 methylation (15, 16). Our results indicate that in plants, GCN5 plays this role. However, transcriptional de-repression is not completely abolished in *gcn5* mutants (Figure 1C), thus suggesting that at least one of the five p300/CBP orthologs in Arabidopsis (HAC1/2/4/5/12 (32, 49)) may also contribute to higher histone acetylation levels in the absence of H3.1K27me1.

Our work shows that GCN5-catalyzed histone acetylation plays a key role in mediating transcriptional activation in *atxr5/6* mutants. The role of GCN5 as a transcriptional co-activator in other biological systems is well defined, thus supporting a conserved function for GCN5 in all eukaryotes. In mammals, H3K27ac is found at active transcriptional enhancers (50, 51), but a recent study in mouse ESCs showed that H3K27ac depletion at enhancers does not affect gene expression (52). This suggests that H3K27ac works redundantly or synergistically with other chromatin features to specify active enhancers. In maize and rice, approximately 30% of H3K27ac sites are found in intergenic regions (42, 53), which support a conserved role for H3K27ac in marking enhancers and contributing to their activity in eukaryotes. However, a recent report has shown that only a small subset of H3K27ac sites (0.5%-3%) are located upstream of the TSS in Arabidopsis, which argues for plant species-specific variation in the types of chromatin features defining active enhancers (54). Aside from marking enhancers, H3K27ac has also been found to be enriched close to the TSS of transcriptionally active protein-coding genes in mammals, maize, rice and Arabidopsis (42, 53–55), a result that we confirmed for Arabidopsis in our ChIP-Rx experiments. H3K36ac has also been shown in multiple biological systems to co-localize with H3K27ac at the TSS of transcriptionally-active regions of the genome (41, 55). These observations suggest important roles for TSS-localized H3K27ac and H3K36ac in mediating transcriptional activity. Precisely mapping the H3K27ac and H3K36ac regions in the heterochromatin of *atxr5/6* mutants in relation to the TSS of de-repressed TEs is challenging, as TSSs are not well defined for TEs. Nevertheless, we did observe H3K27ac and H3K36ac peaks in *atxr5/6* at the 5’end of annotated TEs (Figure 5C, Supplemental Figure 8), which would support a similar mode of action for H3K27ac/H3K36ac in regulating transcription of genes and TEs.

Yeast and animal GCN5 have been shown to have the ability to acetylate multiple lysines (K9, K14, K18, K23, K27 and K36) in the N-terminal tail of histone H3 (18, 20). Our *in vitro* results using recombinant nucleosomes suggest that the GCN5 ortholog in Arabidopsis also has broad substrate specificity. However, the specificity of ATXR5/6 for H3K27 and results from this study suggest a critical role for K27 over other target sites of GCN5 on H3.1 variants. One observation supporting a unique role for H3.1K27ac over other acetylated lysines of H3 in Arabidopsis comes from experiments showing that increased levels of cytosolic acetyl-CoA (the essential cofactor for protein acetylation) increase H3 acetylation in plants (17). Results from these experiments show that H3K27 is predominantly acetylated over other lysine residues of H3 (i.e. H3K9, H3K14 and H3K18; H3K36 was not assessed in that study), in a manner dependent on GCN5. Higher levels of H3K27ac are observed in genic regions, and this correlates with higher transcriptional levels for genes showing gains in H3K27ac (17). Similarly to H3.1K27ac, our *in vitro* and *in vivo* results implicate H3.1K36ac as playing a key role in mediating the heterochromatin phenotypes of *atxr5/6.* However, these results do not rule out the possibility that other acetylated sites (e.g. K9, K14, K18 and K23) on H3.1 also contribute to mediating transcriptional de-repression and genomic instability in plants, for example by acting in a functionally redundant manner. Our *in vitro* histone acetyltransferase assays indicate that deposition of H3K36ac by GCN5 is negatively regulated by H3.1K27me1, although the molecular mechanism responsible for this crosstalk remains unknown. Previous structural work characterizing a protein complex composed of the histone acetyltransferase (HAT) domain of GCN5 from *Tetrahymena thermophila* and a phosphorylated histone H3 peptide (aa. 5-23) showed that the HAT domain interacts with the sidechain of glutamine 5 (Q5), located 9 amino acids upstream of the target lysine (K14) on the H3 peptide (56). As H3K27 is similarly located 9 amino acids upstream H3K36, this suggests that the HAT domain of GCN5 in Arabidopsis may interact with the sidechain of H3K27 to regulate the catalytic activity of GCN5 at H3K36. Structural studies of the HAT domain of Arabidopsis GCN5 will be needed to validate this model.

The catalytic specificity of ATXR5/6 for replication-dependent H3.1 variants together with the observation that heterochromatin amplification is suppressed when the H3.1 chaperone CAF-1 is mutated have led to a model where the H3.1 variant plays a specific role in maintaining genome stability (8). One mechanism that could explain the requirement for H3.1 variants to induce the *atxr5/6* mutant phenotypes would be if GCN5, similarly to ATXR5/6, specifically acetylates K27 in H3.1 variants. However, our results show no difference in enzymatic activity for GCN5 on H3.1 and H3.3 variants (Figure 3A). Therefore, GCN5 is unlikely to be directly involved in mediating the H3.1 requirement for inducing the *atxr5/6* mutant phenotypes. An alternative mechanism to explain the role for H3.1 variants in this process could be that downstream chromatin readers mediating transcriptional de-repression and heterochromatin amplification interact with H3.1K27ac and/or H3.1K36ac, but not H3.3K27ac and/or H3.3K36ac. Another possibility is that transcriptional derepression mediated through GCN5 is not dependent on H3.1 variants, but heterochromatin amplification is. Previous results have shown that expression of ATXR5/6-resistant H3.1A31T (which partially mimics the N-terminal tail of H3.3 variants) in plants generates very low-level transcriptional de-repression in heterochromatin (which is supported by GCN5 being active on H3.3 variants), but genomic instability in the H3.1A31T lines is not detected (8). Therefore, heterochromatin defects in *atxr5/6* mutants could be due to H3.1-independent transcriptional derepression mediated by GCN5-catalyzed H3K27ac and H3K36ac, coupled to another H3.1 variantspecific process that would lead to even higher levels of transcriptional de-repression and heterochromatin amplification. In this two-step model, transcriptional de-repression in heterochromatin via GCN5 in the absence of H3K27me1 would be the initial trigger leading to H3.1-dependent genomic instability. More work will be needed to fully understand the relationship between H3 variants, transcriptional de-repression, and genomic instability.

## Materials and Methods

### Plant materials

Arabidopsis plants were grown under cool-white fluorescent lights (approximately 100 μmol m^-2^ s^-1^) in long-day conditions (16 h light/8 h dark). The *atxr5/6* double mutant was described previously (3). *gcn5 (At3g54610,* SALK_030913), *ada2b (At4g16420,* SALK_019407), *ada3 (At4g29790,* SALK_042026C), *sgf11 (At5g58575,* SAIL_856_F11), *eny2 (At3g27100,* SALK_045015C), *ubp22 (At5g10790,* GK-263H06), *chr5 (At2g13370,* SAIL_504_D01) and *chr6 (At2g25170,* GK-273E06) are in the Col-0 genetic background and were obtained from the Arabidopsis Biological Resource Center (Columbus, Ohio). Temperature-optimized CRISPR/Cas9 was used to generate additional mutant alleles of *GCN5* (in Col-0 and *atxr5/6)* used in this study (23). The *h3.1* quadruple mutant was described previously (8). Transgenic plants expressing WT H3.1 *(At5g65360),* H3.1 S28A, H3.1K9R, H3.1S28A K9R, H3.1K14R, H3.1S28A K14R, H3.1K18R, H3.1S28A K18R, H3.1K23R, H3.1S28A K23R, H3.1K36R, H3.1S28A K36R were made by transforming the *h3.1* quadruple mutant background.

### Constructs

Cloning of the catalytic fragment of *ATXR6* (a.a. 25-349) and the plant PRC2 complexes for protein expression and *in vitro* methyltransferase assays was described previously (3, 8). The histone H3.1 gene *(At5g65360)* and its promoter (1167 bp upstream of the start codon) were cloned into pENTR/D-TOPO (ThermoFisher Scientific, Waltham, MA) and then sub-cloned using Gateway Technology into the plant binary vectors pB7WG (57). Site-directed mutagenesis to generate the different H3.1 point mutant constructs was performed using QuikChange II XL Site-Directed Mutagenesis Kit (Agilent Technologies, Santa Clara, CA). The *ADA2b* coding sequence was cloned into pETDuet-1 (Millipore, Burlington, MA) vector using the SalI and NotI restriction sites, yielding pETDuet-1-ADA2b. The *GCN5* coding sequence was cloned into pETDuet-1-*ADA2b* plasmid using the EcoRV and PacI restriction sites, yielding pETDuet-1-*ADA2b*-*GCN5*. The cloning procedure used to make the CRISPR construct targeting *GCN5* in Arabidopsis was performed as described previously (58).

### Protein expression and purification

Expression and purification of the ATXR6 protein and the plant PRC2 complexes CURLY LEAF and MEDEA has been described previously (3, 8). For the GCN5-ADA2b protein complex, pETDuet-1-ADA2b-GCN5 was transformed into BL21 (DE3) *E. coli* (Millipore), cultured in LB and induced to express proteins by adding 1 mM IPTG. Cells were then pelleted by centrifugation, resuspended in NPI-10 buffer (50mM NaH_2_PO_4_, 300mM NaCl, 10mM Imidazole, pH 8), and lysed by sonification. After centrifugation to remove cell debris, Ni-NTA agarose (Qiagen, Hilden, Germany) was added to the supernatant and rotated at 4 °C for 2 hours. The Ni-NTA agarose was washed 3 times using NPI-20 buffer (50mM NaH_2_PO_4_, 300mM NaCl, 20mM imidazole, pH 8), and the protein complex was eluted in NPI-250 buffer (50mM NaH_2_PO_4_, 300mM NaCl, 250mM imidazole, pH 8). The buffer was changed to 1×PBS (137 mM NaCl, 10 mM phosphate, 2.7 mM KCl, pH 7.4) containing 10% glycerol using an Amicon Ultra-0.5 Centrifugal Filter Unit (30kDa cutoff). The proteins were aliquoted, flash-frozen in liquid nitrogen, and then stored at −80 °C.

The protocols to generate the H3K27me1 and H3K36me2 methyl-lysine analog-containing histones and to make the recombinant chromatin used in the *in vitro* histone modification assays (methylation and acetylation) was described previously (45).

### Histone lysine methyltransferase (HMT) and acetyltransferase (HAT) assays

The general procedure used to perform the *in vitro* histone modification assays presented in this study were described in detail in a previous publication (59).

For the radioactive HMT assays, 0.5 μg of ATXR6, 1.5 μg of MEA or 1.5 μg of CLF (PRC2) complexes were incubated with 1 μg of Histone H3 peptides (GenScript, Piscataway, NJ) and 1.5 μCi of ^3^H-SAM (Perkin Elmer, Waltham, MA) in a 25 μl reaction. The histone methyltransferase buffer contained 50 mM Tris pH 8.0, 2.5 mM MgCl_2_ and 4 mM DTT. The methylation reactions were incubated at 22°C for 2 hours. The samples were pipetted onto Whatman P-81 filter paper and dried for 15 minutes. The free ^3^H-SAM was removed by washing 3 x 30 minutes in 50 mM NaHCO_3_ pH 9.0. The filter paper was dried and added to a vial containing Opti-Fluor^®^ O (Perkin Elmer). Radioactivity on the filter papers was determined using a liquid scintillation counter (Perkin Elmer).

For the HAT assays with antibody detection, 1 μg of recombinant nucleosomes and 2 μg of the GCN5-ADA2b complex were incubated in 50 μl histone acetyltransferase (HAT) buffer (1 mM HEPES pH 7.3, 0.02% BSA) containing 50 mM acetyl co-enzyme A (Acetyl-CoA; Sigma) at 23 °C for 3 hours (wild type H3.1, H3.1K27M and H3.3 nucleosomes) or 5 hours (H3K27me0, H3K27me1, H3K36me0 and H3K36me3 nucleosomes). The reactions were stopped by adding 4X Laemmli Sample Buffer (Bio-Rad) and boiling at 95 °C for 5 min. The samples were resolved by 15% SDS-PAGE gel, transferred to PVDF membrane, and western blot was performed using anti-H3K9ac (Cell Signaling Technology, Danvers, MA: 9649), anti-H3K14ac (Active Motif, Carlsbad, CA: 39698), anti-H3K18ac (Active Motif: 39588), anti-H3K23ac (Active Motif: 39132), anti-H3K27ac (Active Motif: 39135), anti-H3K36ac (Active Motif: 39379) or anti-H3 antibodies (Abcam: ab1791) and a secondary anti-Rabbit HRP-labeled antibody (Sigma).

For the radioactive HAT assays, 1 μg of peptides and 1 μg of GCN5-ADA2 complex were incubated in 25 μl HAT buffer containing 0.625 μCi ^3^H-Acetyl-CoA (PerkinElmer) at 23 °C for 2 hours. Reactions were stopped by pipetting onto Whatman P-81 filter paper and activity (cpm) was measured using a liquid scintillation counter (Perkin Elmer).

### Chromatin Immunoprecipitation

ChIP was performed as described previously (60), with some modifications. Leaves from three-week-old plants were fixed in 1% formaldehyde. Immunoprecipitation was performed using protein A magnetic beads (New England BioLabs, Ipswich, MA). Following the Proteinase K treatment of each sample, immunoprecipitated DNA was purified using ChIP DNA Clean & Concentrator kit (Zymo Research, Irvine, CA). 2 μl of Histone H3 antibody (Abcam: ab1791), 2.5 μl of H3K27ac antibody (Active Motif: 39135), 5 μl of H3K36ac antibody (Active Motif: 39379) or 2.5 μl of H3K36me3 (Abcam: ab9050), was used per immunoprecipitation (750 μl of chromatin solution). For the H3K27ac and H3K36ac ChIP experiments, ChIP with exogenous genome (ChIP-Rx) was performed in order to properly normalize the data (40). For each sample, an equal amount of drosophila chromatin (Active Motif: 53083) was added prior to chromatin shearing.

### Nuclei DAPI staining

Leaves from four-week-old plants were fixed in 3.7% formaldehyde in cold Tris buffer (10 mM Tris-HCl pH 7.5, 10 mM NaEDTA, 100 mM NaCl) for 20 minutes. Formaldehyde solution was removed, and leaves were washed twice for 10 minutes in Tris buffer. The leaves were then finely chopped with razor blade in 500 μl LB01 buffer (15 mM Tris-HCl pH7.5, 2 mM NaEDTA, 0.5 mM spermine-4HCl, 80 mM KCl, 20 mM NaCl and 0.1% Triton X-100). The lysate was filtered through a 30 μm mesh (Sysmex Partec, Gorlitz, Germany). 5 μl of lysate was added to 10 μl of sorting buffer (100 mM Tris-HCl pH 7.5, 50 mM KCl, 2mM MgCl_2_, 0.05% Tween-20 and 5% sucrose) and spread onto a coverslip until dried. Cold methanol was added onto each coverslip for 3 min, then rehydrated with TBS-Tx (20 mM Tris pH 7.5, 100 mM NaCl, 0.1% Triton X-100) for 5 min. The coverslips were mounted onto slides with Vectashield mounting medium DAPI (Vector Laboratories, Burlingame, CA). Nuclei were imaged on a Nikon Eclipse Ni-E microscope with a 100X CFI PlanApo Lamda objective (Nikon, Minato City, Tokyo, Japan). Digital images were obtained using an Andor Clara camera. Z-series optical sections of each nucleus were obtained at 0.3 μm steps. Images were deconvolved by imageJ using the deconvolution plugin.

### RT-qPCR

Total RNA was extracted from three-week-old leaf tissue using TRIzol (Invitrogen, Carlsbad, CA). Samples were treated with RQ1 RNase-free DNase (Promega, Madison, WI) at 37°C for 30 min. SuperScript II Reverse Transcriptase (Invitrogen) was used to produce cDNA from 1 μg of total RNA. Reverse transcription was initiated using oligo dT primers. Quantification of cDNA was done by realtime PCR using a CFX96 Real-Time PCR Detection System (Bio-Rad, Hercules, CA) with KAPA SYBR FAST qPCR Master Mix (2×) Kit (Kapa Biosystems, Wilmington, MA). Each primer pair was assessed for efficiency of amplification. Relative quantities were determined by the Ct method (61). Actin was used as the normalizer. At least three biological samples were used for each experiment.

### Flow cytometry

Rosette leaves from three-week-old plants were finely chopped in 0.5 ml Galbraith buffer (45 mM MgCl_2_, 20 mM MOPS, 30 mM sodium citrate, 0.1% Triton X-100, 40 μg/μl RNase A) using a razor blade. The lysate was filtered through a 30 μm mesh (Sysmex Partec). Propidium iodide (Sigma, St. Louis, MO) was added to each sample to a concentration of 20 μg/ml and vortexed for 3 seconds. Each sample was analyzed using a BD FACS LSR Fortessa X20 (Becton Dickinson, Franklin Lakes, NJ). Quantification (nuclei counts and robust CV values) was performed using Flowjo 10.0.6 (Tree Star, Ashland, Oregon).

### Next-generation sequencing library preparation

RNA samples were prepared from three-week old leaf tissue using the RNeasy Plant Mini Kit (Qiagen). RNA and ChIP sequencing libraries were prepared at the Yale Center for Genome Analysis (YCGA). RNA samples were quantified and checked for quality using the Agilent 2100 Bioanalyzer Nano RNA Assay. Library preparation was performed using Illumina’s TruSeq Stranded Total RNA with Ribo-Zero Plant in which samples were normalized with a total RNA input of 1 μg and library amplification with 8 PCR cycles. ChIP library preparation was performed using TruSeq Library Prep Kit (Illumina, San Diego, CA). Libraries were validated using Agilent Bioanalyzer 2100 High sensitivity DNA assay and quantified using the KAPA Library Quantification Kit for Illumina^®^ Platforms kit. Sequencing was done on an Illumina NovaSeq 6000 using the S4 XP workflow.

### RNA-seq processing and analysis

Two independent biological replicates for Col, *atxr5/6*, *gcn5* and *atxr5/6 gcn5* were sequenced. Paired-end reads were filtered and trimmed using BBTools (version 38.79) (62). Reads with quality inferior to 20 were removed. The resulting data sets were aligned against the *Arabidopis* genome (TAIR10) using STAR (version 2.7.2a) allowing 2 mismatches (--outFilterMismatchNmax 2) (63). Consistency between biological replicates was confirmed by Pearson correlation using deepTools2 (Supplementary Figure 9) (64). Protein-coding genes and transposable elements (TE) were defined as described in the TAIR10 annotation gff3 file. The program featureCounts (version 1.6.4) (65) was used to count the paired-end fragments overlapping with the annotated protein-coding genes and TEs. Differential expression analysis of protein-coding genes was performed using DESeq2 version 1.26 (66) on raw read counts to obtain normalized fold changes (FC) and *Padj-values* for each gene. Genes were considered to be differentially expressed only if they showed a log2FC >1 and a *Padj-values* < 0.05. TPM (transcripts per million) values were calculated for TEs. To define TEs as up-regulated in the *atxr5/6* mutant, they must show 2-fold up-regulation as compared to Col in both biological replicates and have a value of TPM > 5. The heatmap was drawn with R program (version 3.6.2) (67).

### ChIP-seq processing and analysis

Two independent biological replicates for Col, *atxr5/6*, *gcn5* and *atxr5/6 gcn5* were sequenced. In order to properly compare H3K27ac and H3K36ac levels between each genotype, we performed ChIP with reference exogenous genome (ChIP-Rx) (40) using equal amounts of drosophila chromatin in each sample as reference. Paired-end reads were filtered and trimmed using BBTools (62). Reads with quality inferior to 20 were removed. Data sets were aligned against combined genomes of *Arabidopsis thaliana* (TAIR10) and *Drosophila melanogaster* (dm6) using bowtie2 (68) with default parameters. Duplicate reads were removed using Picard toolkit (69) (MarkesDuplicates with *REMOVE_DUPLICATES=true).* Consistency between biological replicates was confirmed by Pearson correlation using deepTools2 (Supplementary Figure 10) (64). To calculate the Rx scaling factor of each biological replicate, Drosophila-derived IP read counts were normalized according to the number of input reads. Spike-in normalization was performed as previously described (70). The Rx factors are presented in Supplementary Table 4. We generated bedgraph files with a bin size of 10 bp using deepTools. The bedgraph files were then scaled by adjusting the number of reads in each bin with the Rx factors and therefore generating reference-adjusted reads per million (RRPM). H3K27ac and H3K36ac enriched regions were identified by computing the differential between each bin (± 1kb) to define local maxima.

The number of reads corresponding to euchromatic regions was much higher than the ones from heterochromatic regions. To best determine the heterochromatic enrichment of H3K27ac in each genotype of interest, we avoided the “noise” from the euchromatic reads by first defining heterochromatic regions and extracting the corresponding reads from each genotype. We defined the heterochromatic regions based on the chromatin states proposed by Sequeira-Mendes *et al.*, 2014 (43). We attributed the value of the state number (1 to 9) for each bin of the Sequeira-Mendes *et al.* annotation, and averaged them on 100 kb windows. Only the 100 kb windows with a score superior to 7 were considered as heterochromatic regions (Supplementary Table 2). We then generated a bam file with the reads corresponding to the defined heterochromatic regions. We identified heterochromatic H3K27ac an H3K36ac enriched regions by calculating the log2 ratio between H3K27ac or H3K36ac IP and H3 input using the heterochromatin bam file. The enriched regions were defined with the following criteria: log2 (IP/H3) > 0.3. To compare the H3K27ac and H3K36ac enriched regions between Col and our mutant genotypes, we computed log2 (mutant/Col), using the Rx factor normalized bedgraph file. We considered the levels of H3K27ac and H3K36ac to be differential between genotypes when log2 (mutant/Col) > 0.8. These regions needed to be detected in both replicate in order to be considered.

### Primers

All primers used in this study are listed in Supplemental Table 7.

### Data availability

Sequencing data are available at the Gene Expression Omnibus (GEO) under accession code GSE146126.

## Supplemental Figures

**Supplemental Figure 1.**
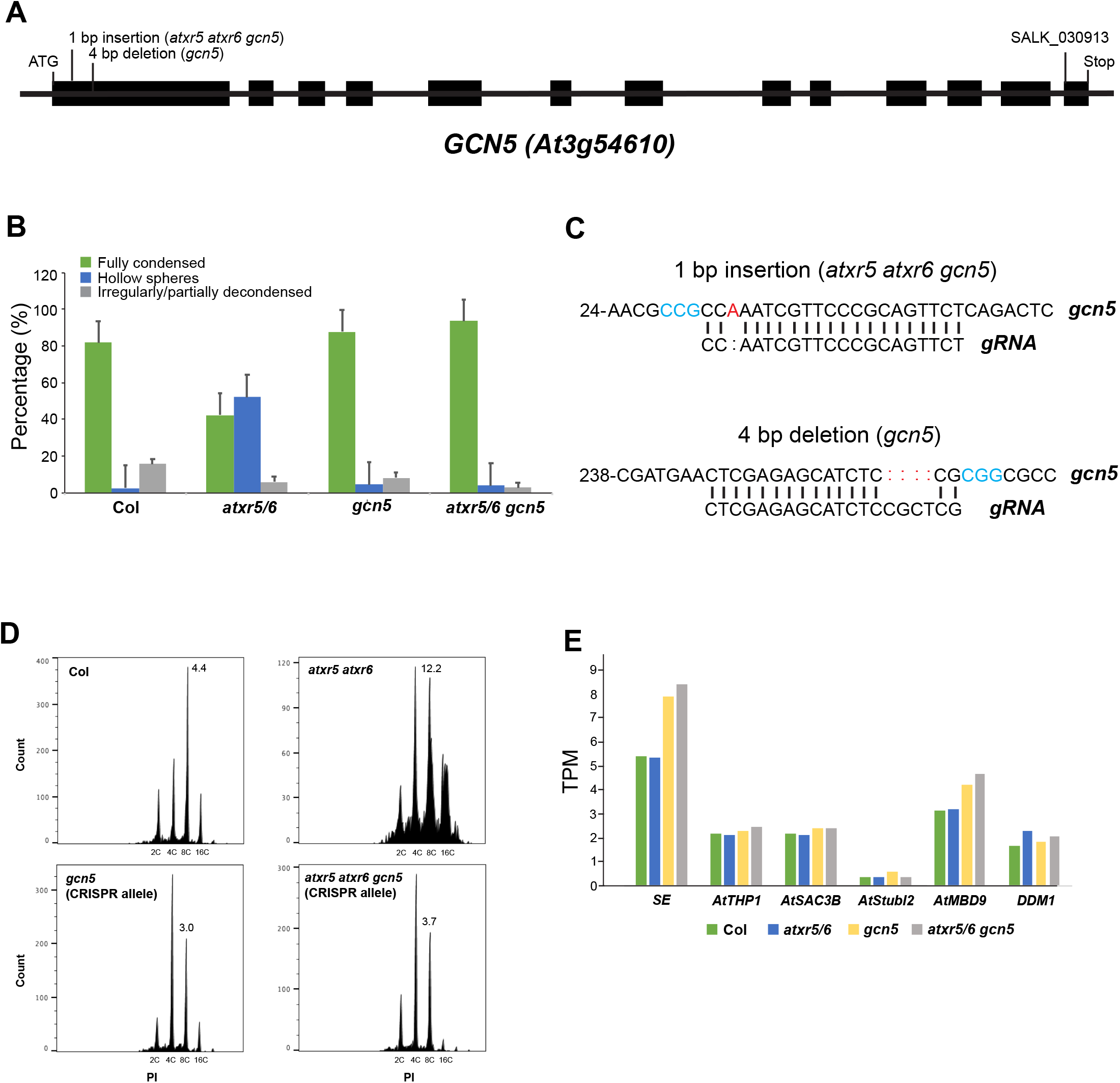
Effect of *GCN5* on genome stability and transcriptional de-repression. (A) Gene structure for *GCN5.* Exons are highlighted as black boxes. The location of the mutations in the *gcn5* alleles used in this study are shown. (B) Quantification of chromocenter appearance. Shown is the percentage of nuclei that are fully condensed (green), hollow spheres characteristic of the *atxr5/6* mutant plants (blue) and irregularly/partially decondensed (grey). At least 25 nuclei for 3 biological replicates of each genotype were assessed. Error bars indicate SEM (C) CRISPR-induced mutations of *GCN5* in Col and *atxr5/6* backgrounds. Mutations (red) and PAM motif (blue) are shown. (D) Flow cytometry profiles of Col, *atxr5/6* and the CRISPR-induced knockout allele of *gcn5* in Col and the *atxr5/6* mutant background. The numbers below the peaks indicate ploidy levels of the nuclei. The numbers above the 16C peaks indicate the robust CV. (E) Gene expression levels in our RNA-seq experiments for the known suppressors of transcriptional de-repression in *atxr5/6.*

**Supplemental Figure 2.**
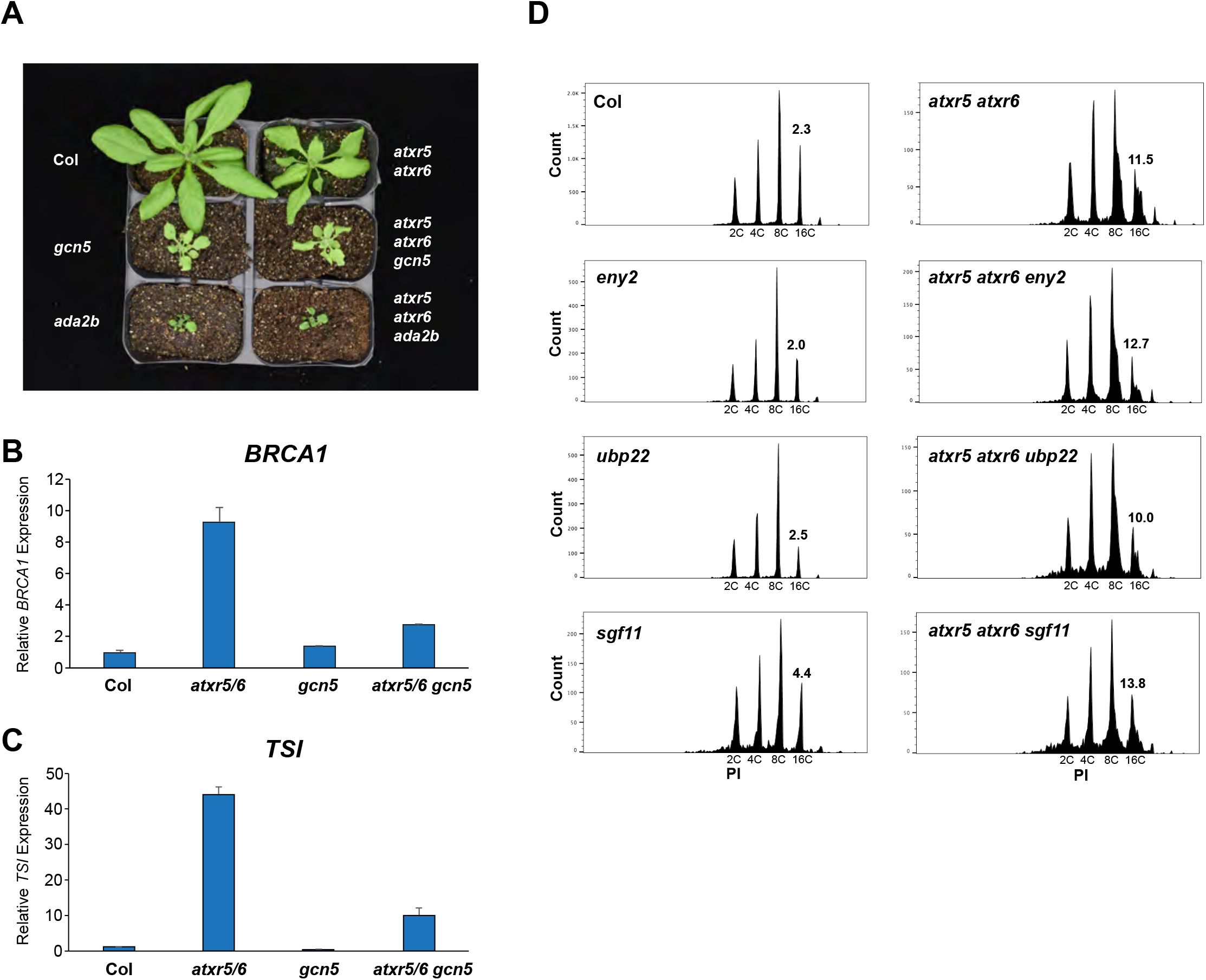
Role of SAGA-related proteins in transcriptional de-repression and genome stability. (A) Growth phenotype of *gcn5* and *ada2b* single mutants and in combination with *atxr5/6*. (B and C) RT-qPCR for the genome stability marker *BRCA1* (B) and the heterochromatic transcriptional reactivation marker *TSI* (C). The average of three biological replicates and SEM are shown. (D) Flow cytometry profiles of the mutant alleles of genes predicted to code for subunits of SAGA in plants. The Robust CV value calculated for the 16C peak on each plot is used as a measure of heterochromatin amplification.

**Supplemental Figure 3.**
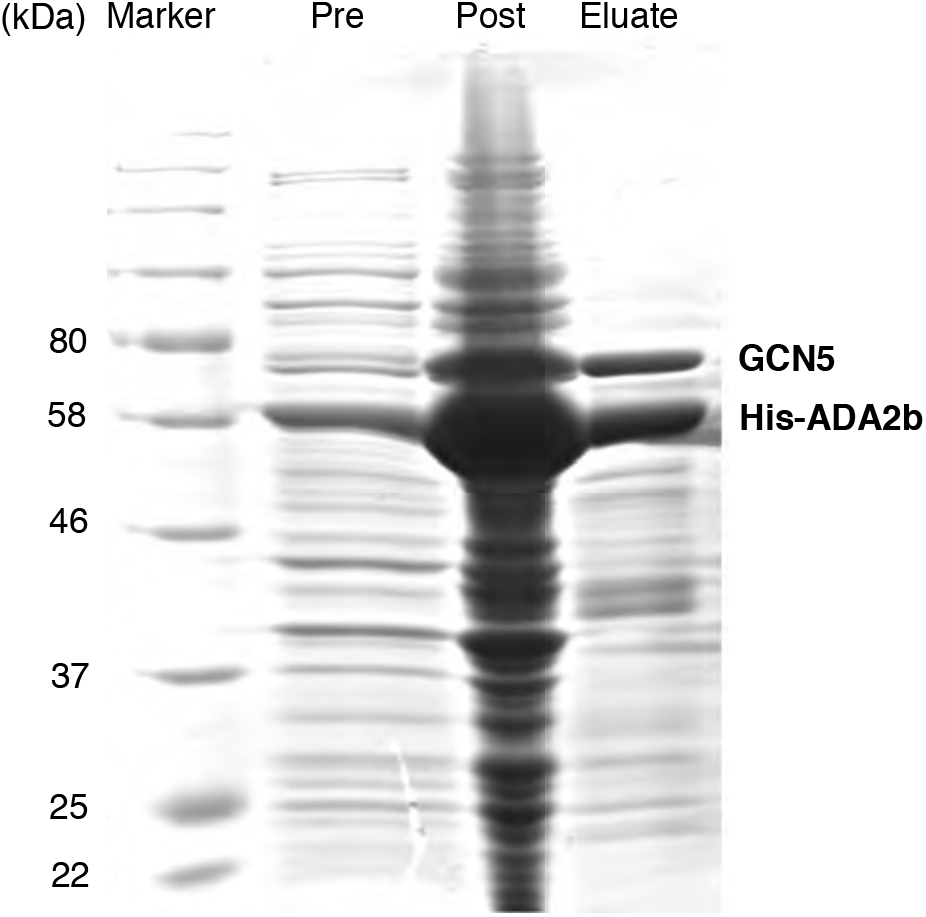
Purification of the GCN5-ADA2b complex. Coomassie-stained gel showing GCN5-ADA2b protein expression either pre- or post-induction in *E. coli,* or after affinity purification (eluate).

**Supplemental Figure 4.**
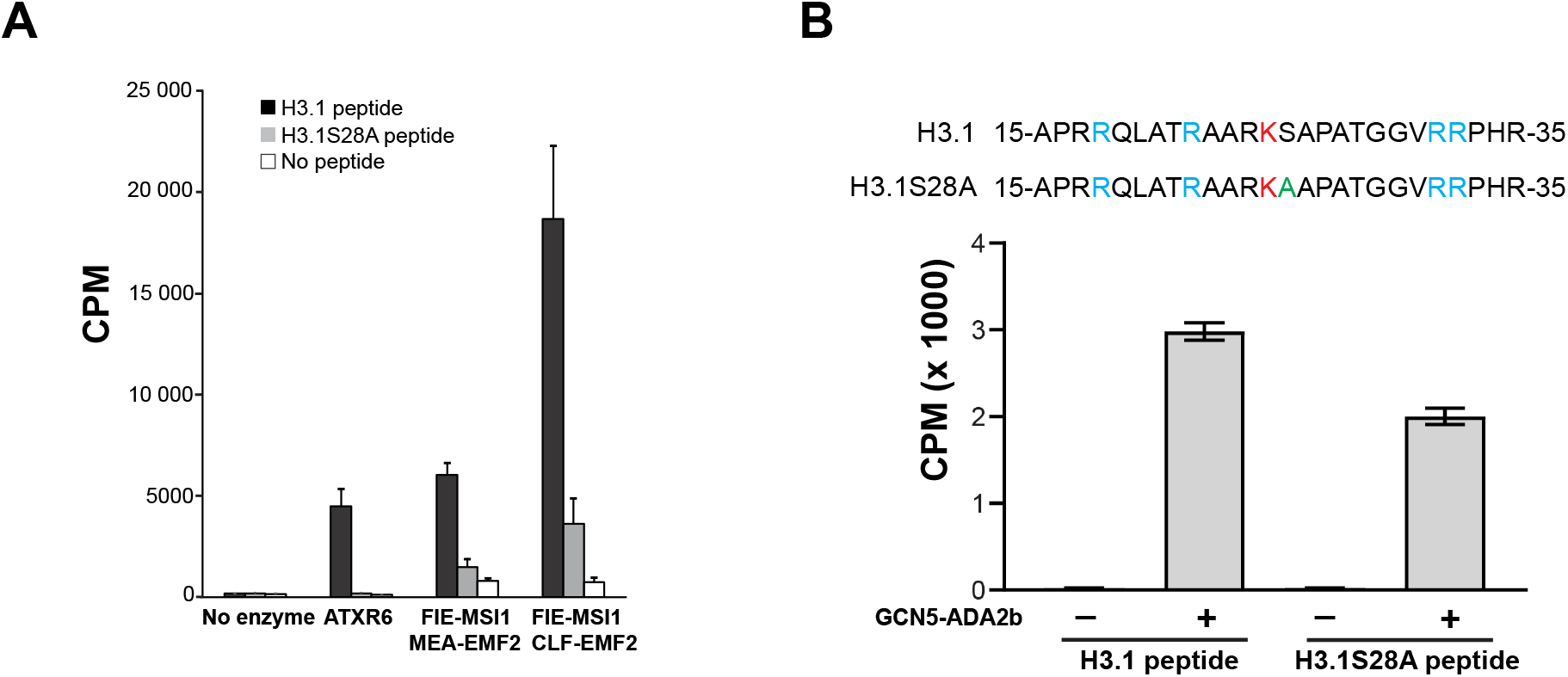
*In vitro* histone modification assays. (A) *In vitro* histone lysine methylation assays at H3.1K27 using peptide substrates, ATXR6 and plant PRC2 complexes. The average of three experiments and SEM are shown. (B) *In vitro* HAT assays using H3.1 peptides. Lysine (K) to arginine (R) mutations were introduced (blue) on the peptides at other potential targets (H3.1K18, H3.1K23, H3.1K36 and H3.1K37) of GCN5-ADA2b, so that the acetylation signal could be specifically measured at H3.1K27 (red). H3.1S28A is shown in green. The average of three experiments and SEM are shown.

**Supplemental Figure 5.**
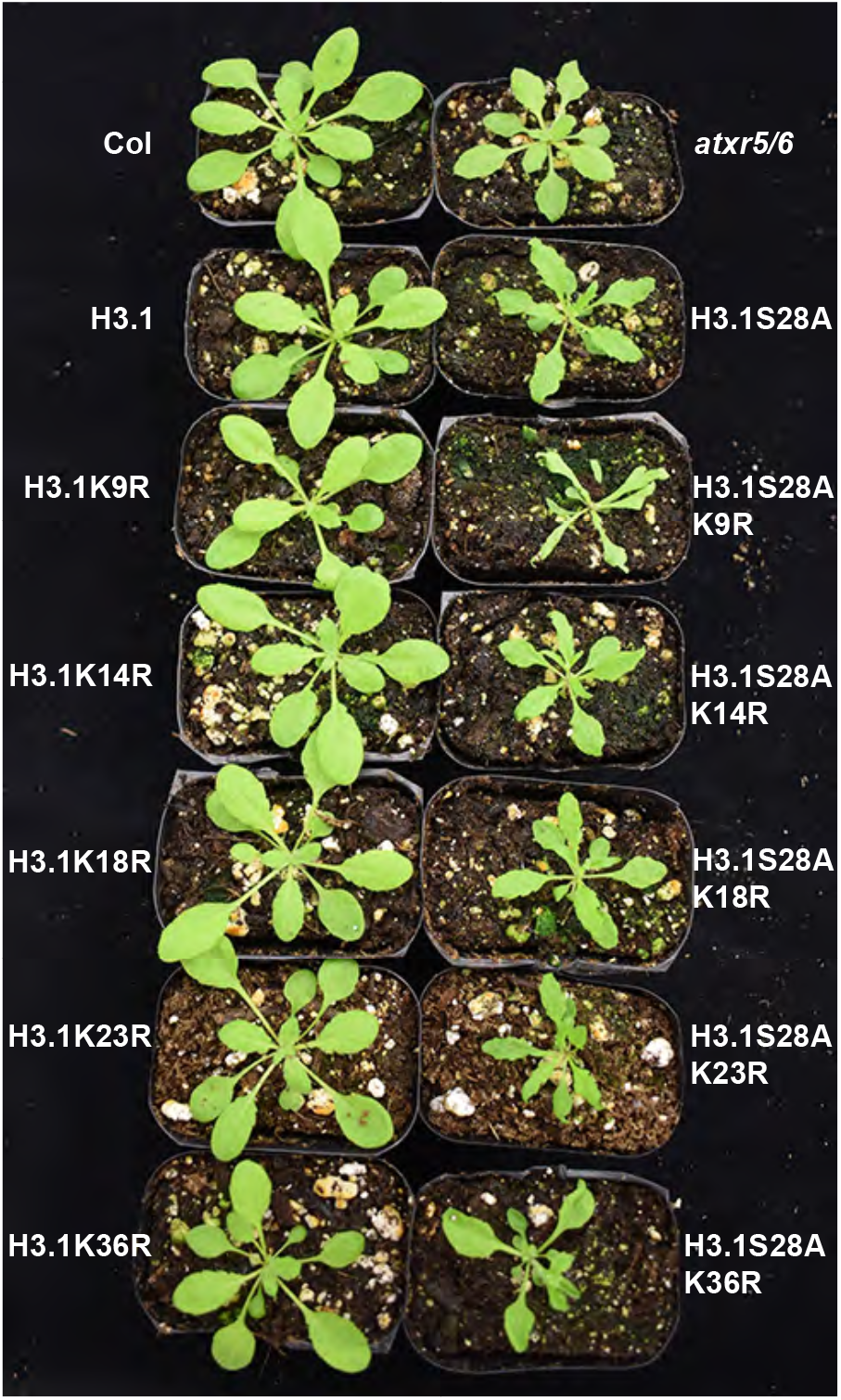
Growth and developmental phenotypes of T1 plants expressing different H3.1 transgenes.

**Supplemental Figure 6.**
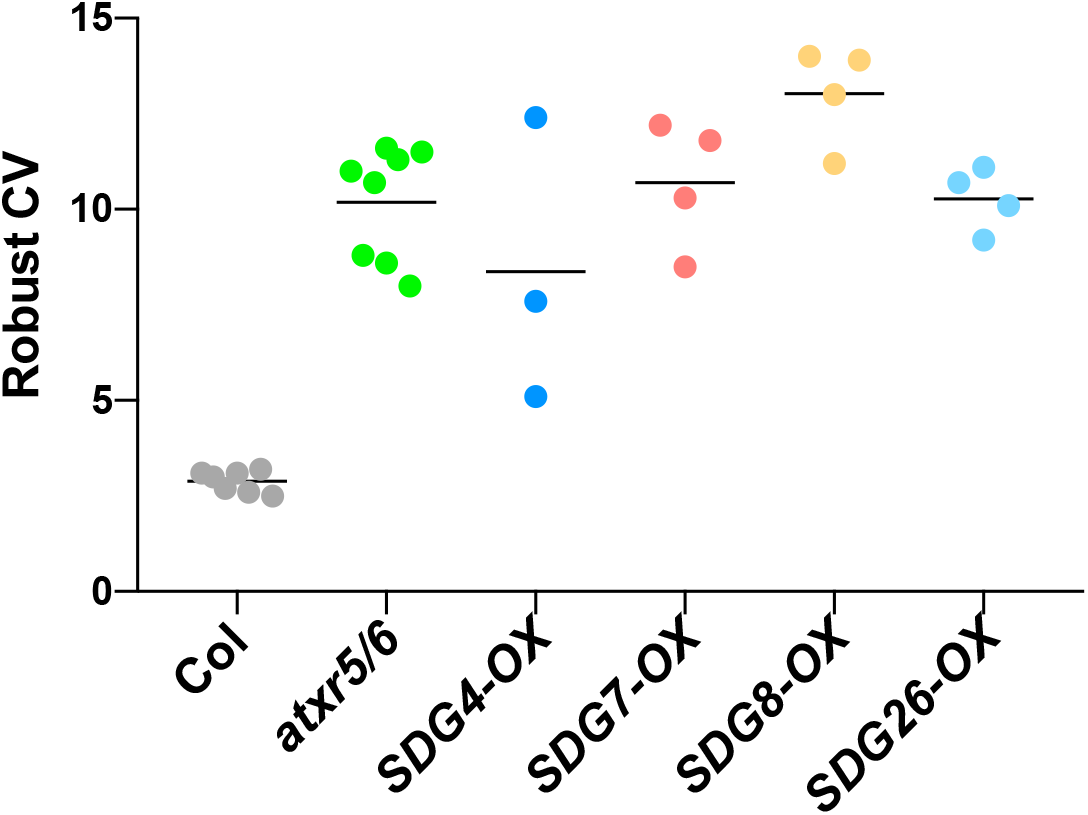
Robust CV values for 16N nuclei obtained by flow cytometry analyses. For Col and *atxr5/6,* each dot represents an independent biological replicate. For overexpression lines, each dot represents one first-generation transformed (T1) plant.

**Supplemental Figure 7.**
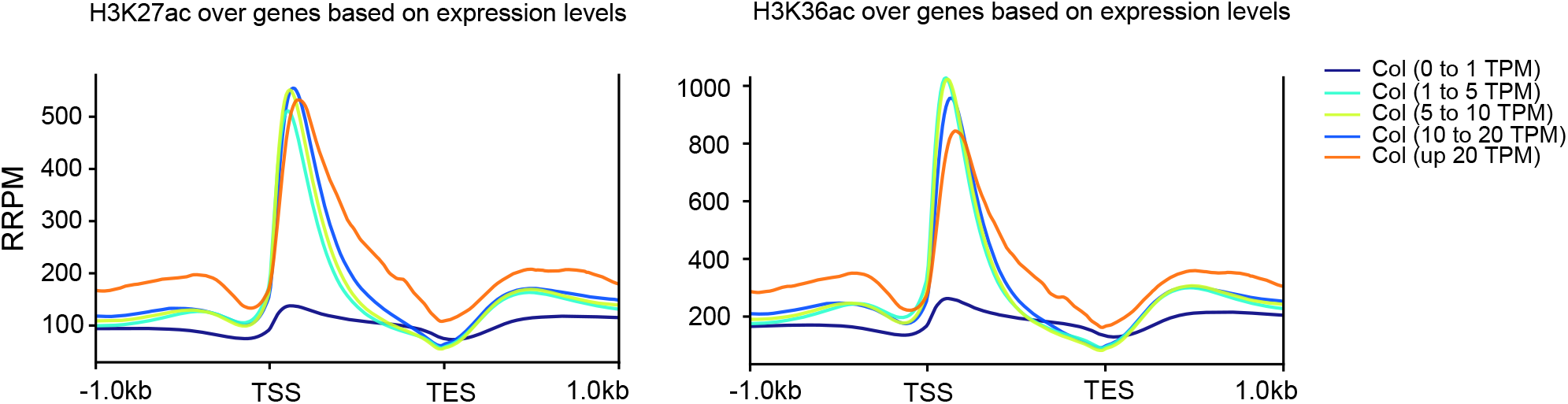
Average distribution of H3K27ac and H3K36ac over protein-coding genes grouped by their expression levels. TSS, transcription start site; TES, transcription end site.

**Supplemental Figure 8.**
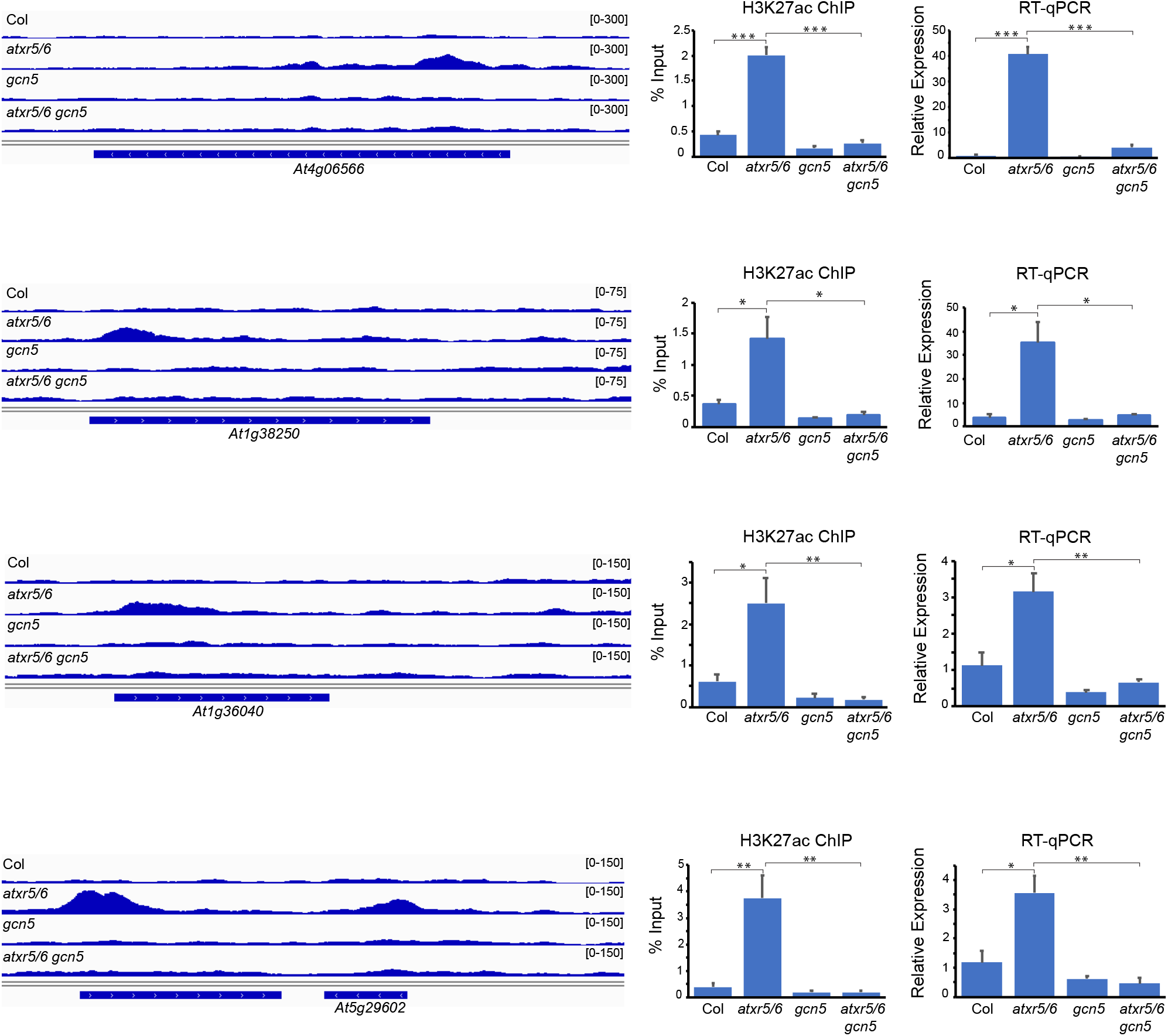
Validation of ChIP-seq and RNA-seq analyses. Genome browser snapshots at different TEs showing H3K27ac enrichment in *atxr5/6*, ChIP-qPCR confirmation of H3K27ac enrichment and expression levels for these TEs. *At4g06566* and *At1g38250* were detected as de-repressed in *atxr5/6* by RNA-seq, but not *At1g36040* or *At5g29602*. Data represents the mean of three biological replicates and error bars indicate SEM. Unpaired t-test: * *p* < 0.05, ** *p* < 0.01 and \*\*\**p* < 0.001.

**Supplemental Figure 9.**
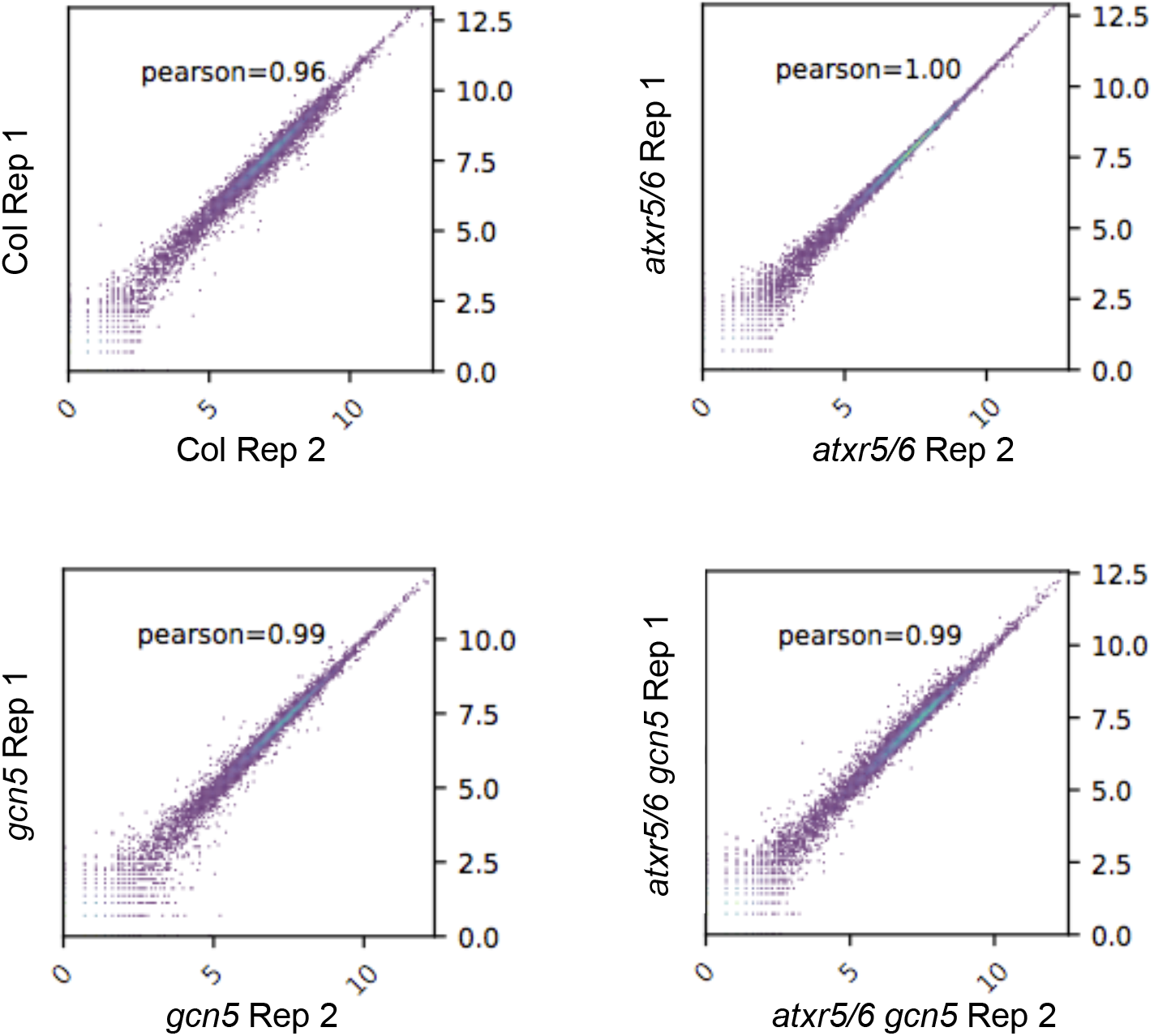
Scatterplots and Pearson correlation coefficients for RNA-seq replicates of Col, *atxr5/6, gcn5* and *atxr5/6 gcn5.*

**Supplemental Figure 10.**
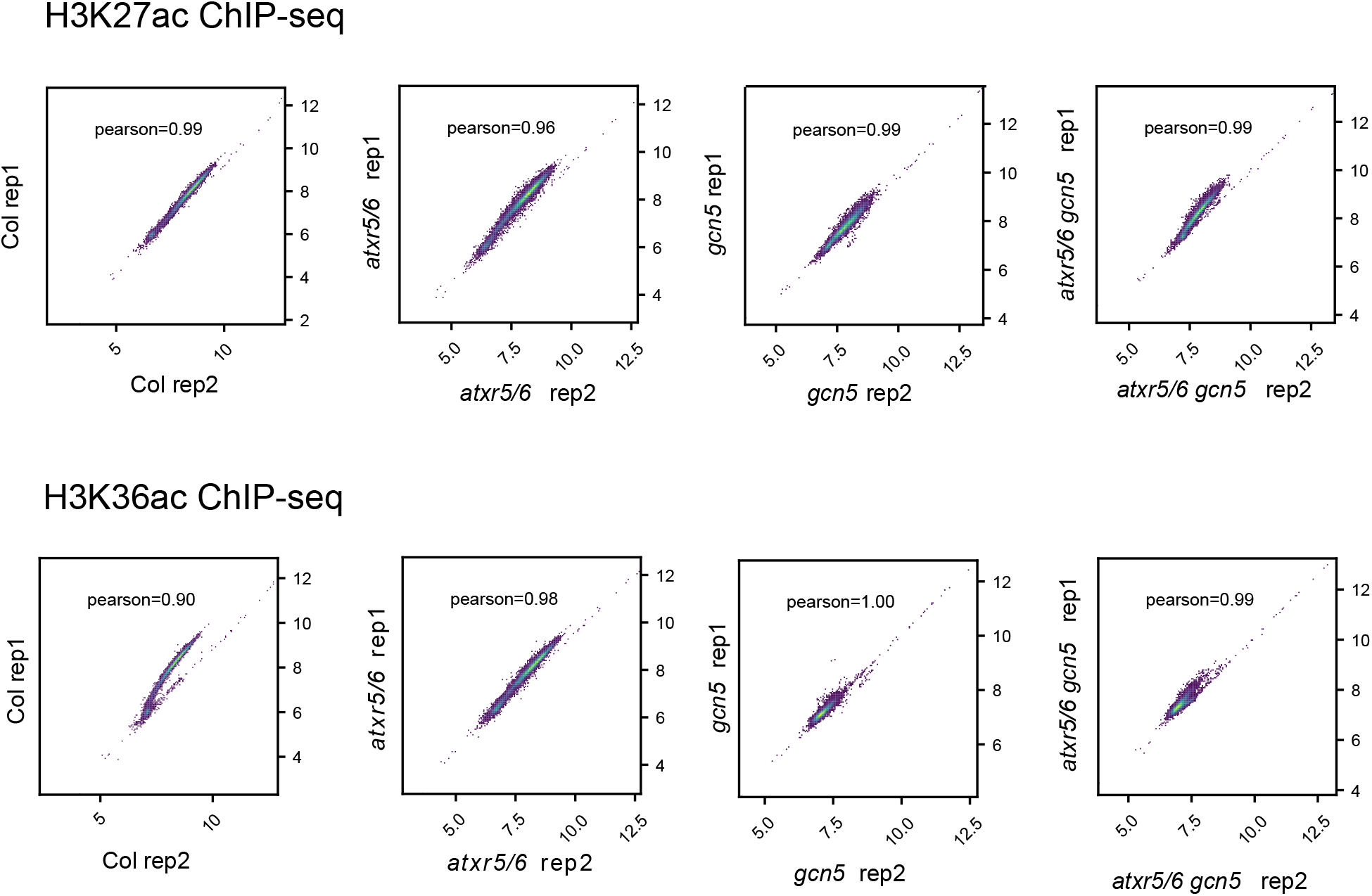
Scatterplots and Pearson correlation coefficients for H3K27ac and H3K36ac ChIP-seq replicates of Col, *atxr5/6, gcn5* and *atxr5/6 gcn5.*

## Supplementary Tables

**Supplemental Table 1. TEs de-repressed in *atxr5/6.*** TEs highlighted in blue are detected only at the highest sequencing depth. TEs in green are detected only at the lowest sequencing depth. TEs highlighted in gray are detected at the highest and lowest sequencing depths.

**Supplemental Table 2. Misregulated genes in *atxr5/6*, *gcn5* and *atxr5/6 gcn5*.**

**Supplemental Table 3. Regions of Arabidopsis genome defined as heterochromatin.**

**Supplemental Table 4. Heterochromatic regions enriched in H3K27ac and H3K36ac in *atxr5/6.***

**Supplemental Table 5. TEs that are de-repressed and overlap with heterochromatic regions enriched in H3K27ac and H3K36ac in *atxr5/6*.**

**Supplemental Table 6.** Rx factors for Col, *atxr5/6*, *gcn5* and *atxr5/6 gcn5* replicates.

|  | **Col**<br>**Rep1** | **Col**<br>**Rep2** | ***atxr5/6***<br>**Rep1** | ***atxr5/6***<br>**Rep2** | ***gcn5***<br>**Rep1** | ***gcn5***<br>**Rep2** | ***atxr5/6***<br>***gcn5***<br>**Rep1** | ***atxr5/6***<br>***gcn5***<br>**Rep2** |
| --- | --- | --- | --- | --- | --- | --- | --- | --- |
| **D.**<br>**melanogaster**<br>**unique H3**<br>**reads** | 51265 | 134411 | 69402 | 104229 | 114932 | 100768 | 69215 | 115431 |
| **A. thaliana**<br>**unique H3**<br>**reads** | 30827170 | 57924375 | 39621511 | 50359868 | 54725117 | 44476014 | 49878156 | 44258118 |
| **% of D.**<br>**melanogaster**<br>**H3 reads** | 0.16629811 | 0.23204566 | 0.17516243 | 0.20696837 | 0.21001691 | 0.22656707 | 0.13876816 | 0.26081317 |
| **D.**<br>**melanogaster**<br>**H3K27ac**<br>**reads** | 73203 | 154549 | 69048 | 88588 | 211097 | 172676 | 161463 | 171638 |
| **A. thaliana**<br>**H3K27ac**<br>**reads** | 35425890 | 53958274 | 39611214 | 38922755 | 34654299 | 36321638 | 50898781 | 35808072 |
| **H3K27ac Rx**<br>**factor** | **2.2717389** | **1.50143749** | **2.53682114** | **2.33630259** | **0.99488344** | **1.31209355** | **0.85944248** | **1.51955376** |
| **D.**<br>**melanogaster**<br>**H3K36ac**<br>**reads** | 100430 | 134550 | 120870 | 126640 | 355181 | 302930 | 35828 | 223401 |
| **A. thaliana**<br>**H3K36ac**<br>**reads** | 40539185 | 47383778 | 38801208 | 41274576 | 20973432 | 19928195 | 25937684 | 18896708 |
| **H3K36ac Rx**<br>**factor** | **1.65586086** | **1.72460545** | **1.44918033** | **1.63430491** | **0.59129545** | **0.74791888** | **0.413212005** | **1.167466434** |

**Supplemental Table 7.**
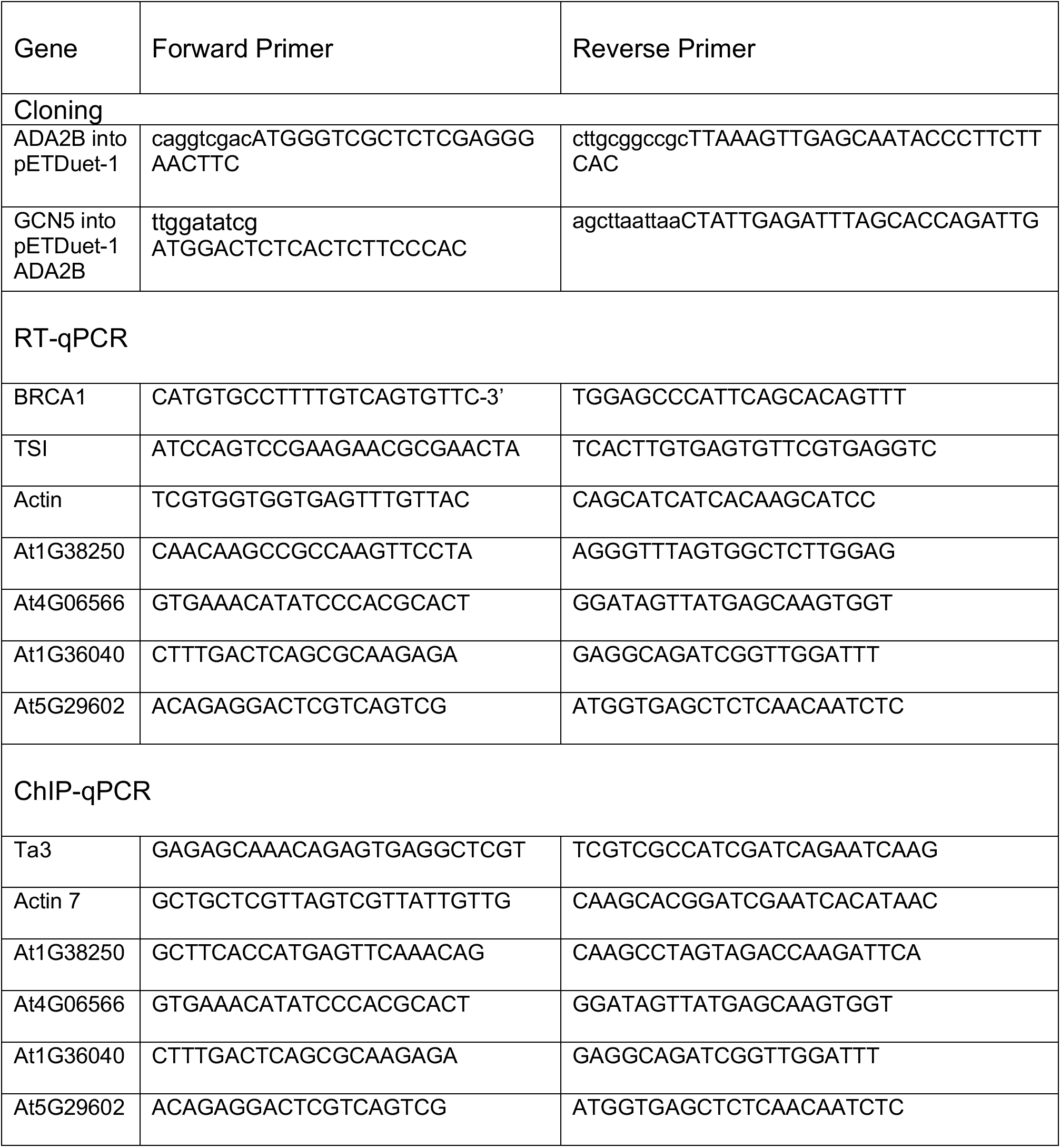
Cloning and PCR primers.

## Author Contributions

Y.J., J.D., C.L. and A.P. designed the experiments. Y.J. wrote the paper with contributions from J.D., C.L. and A.P. All *in vitro* assays were performed by J.D. C.L. performed the ChIP experiments. A.P. did the bioinformatics analyses of all ChIP-seq experiments. A.P. and V.J. did RNA-seq analyses. Microscopy was done by C.L. Flow cytometry analyses were performed by C.L., B.M, G.V and J.M. RNA extractions and RT-qPCR were done by C.L. and J.D and G.V. Crosses were done by G.V. and B.M. Genotyping and plant transformations were performed by G.V., J.M., C.L., and B.M. G.V. made the CRISPR/Cas9 mutants. K.M.W. and P.V. made the modified and unmodified nucleosomes used in the *in vitro* assays.

## Acknowledgments

We thank members of our lab for discussions and advice during the course of this work. We want to acknowledge Christopher Bolick and his staff at Yale for help with plant growth and maintenance. We also thank Jean-François Couture (University of Ottawa) for sending the K27M nucleosomes used in this study, and Kenneth Nelson (Yale University) for technical help with flow cytometry. This project was supported by grant #R35GM128661 from the National Institutes of Health to Y.J. B.M. was supported by a Yale University Brown Fellowship. V.J. is supported by the Fonds de Recherche du Québec-Nature et Technologies (FRQNT) [272565]. Work in the Voigt lab is supported by the Wellcome Trust ([104175/Z/14/Z], Sir Henry Dale Fellowship to P.V.) and the European Research Council (ERC) under the European Union’s Horizon 2020 research and innovation programme (ERC-STG grant agreement No. 639253). The Wellcome Centre for Cell Biology is supported by core funding from the Wellcome Trust [203149]. We are grateful to the Edinburgh Protein Production Facility (EPPF) for their support. The EPPF was supported by the Wellcome Trust through a MultiUser Equipment grant [101527/Z/13/Z]. The authors declare that they have no competing interests.

